# Critical period plasticity enables credit assignment

**DOI:** 10.64898/2026.06.24.734069

**Authors:** Ruth J. Meier, Salomon Muller, Bin Wang, Kamila Mizerska, Sania Jain, Thomas Fothergill, Julia Mercer, Coleman Klapheke, Julia DiSano, Nikola Milicic, Ruohong Wang, Sujatha Narayan, Joe Li, Kurt Weiss, Efrén Álvarez-Salvado, Misha B. Ahrens, Masahiko Hibi, Jan Huisken, Kevin W. Eliceiri, David E. Ehrlich

## Abstract

Synaptic plasticity is often guided by instructive inputs to neural circuits, but learning only succeeds when these instructions reach neurons that mediate relevant outputs. This creates the credit assignment problem: how does a neuron receive instructions suited to its own behavioral function? Here we report a developmental mechanism that enables credit assignment by using instructive inputs to organize the downstream architecture through which learning is expressed. In the olivocerebellar learning system, the inferior olive provides instructive inputs that guide plasticity within the cerebellum. During circuit assembly in zebrafish, we find these same inputs regulate long-range cerebellar projections during a highly plastic, two-day critical period. Developmental experience caused specific maturation of projections to targets that were coactivated with olivary inputs. Mathematical theory and computational modeling show how the resulting architecture constrains later learning, such that a single input can sculpt downstream connectivity and then leverage that circuit to assign credit. Simulated learning became paradoxically more robust if we reduced plasticity after a developmental critical period, protecting key architecture from corruption during learning. Thus, instructive inputs can first build the circuits they later teach, coordinating development and learning to enable effective credit assignment.

## Introduction

Throughout the brain, learning is encoded by plasticity at synapses far upstream of where behavioral consequences ultimately emerge. For synaptic changes to reliably improve performance, instructive inputs that guide plasticity must reach the specific neurons capable of appropriately modifying network output, a challenge known as the credit assignment problem^1,2^. Neural circuits across phylogeny and brain region assign credit by virtue of their architecture, constraining which neurons receive a given instruction and thereby ensuring plasticity is directed to sites of suitable downstream influence^3–5^. For example, precise anatomy organizes cerebellar learning, in which the inferior olive (IO) delivers instructions to the specific Purkinje cells at which local plasticity can improve performance, despite lacking explicit knowledge of the downstream structures through which plasticity manifests behaviorally^6–13^. However, the origins of anatomical mechanisms for credit assignment remain mysterious.

Circuit architecture for credit assignment may be sculpted by experience early in life. While basic circuit blueprints are specified genetically as a scaffold of cell types and projections on which learning will later operate^14–18^, a capacity to adapt wiring to experience may be advantageous or essential. Tasks of high dimensionality and directional specificity require granular registration of instructive inputs^9,15,19^. Furthermore, sensitivity to experience could afford flexibility to adapt circuits to the experience of the organism rather than genetically specified representations and tasks. For example, in sensory systems, where connectivity is commonly adapted to early sensory experience during transient, plastic ‘critical periods’, patterned activity reinforces coactive inputs, prunes inactive ones, and tunes circuits to the statistics of the sensed world^20–22^. Furthermore, early experience regulates not only sensory innervation but also cerebellar projections to the balance-related vestibular nuclei, indicating adaptability of precise cerebellar architecture^23,24^.

Here we examined whether principles of experience-dependent circuit development act meaningfully on credit assignment. Using perturbative experiments in the assembling cerebellum of the externally developing zebrafish combined with mathematical theory and computational modeling, we discovered experience-dependent circuit development acts directly on the architecture for credit assignment. Specifically, we find the instructive pathway from IO regulates Purkinje cell projections logically based on experience, with attendant behavioral changes. We demonstrate how a correlation-based rule for developmental plasticity, guided by the instructive inputs that later guide plasticity for learning, can preconfigure the olivocerebellar system for credit assignment. Finally, we reveal why restricting developmental plasticity to a critical period is essential for supporting lifelong memory expression. Together, these results outline a role for instructive inputs to first construct the circuits they later teach, a single pathway orchestrating development and learning in an economical solution to credit assignment.

## Results

### A critical period for plastic cerebellar projections

We devised an approach to study the role of the IO in cerebellar development and function, leveraging an evolutionarily conserved circuit architecture and the optical transparency of developing zebrafish (**Fig. 1A**)^25^. We screened for developmental plasticity by imaging the assembling olivocerebellar system of 757 fish, enabled by the Flamingo light-sheet microscopy initiative (https://huiskenlab.com/flamingo/). Immature circuits are often sensitive to perturbation only transiently, so we first evaluated developmental plasticity windows. We activated Purkinje cells at variable phases of cerebellar development using a chemogenetic approach. Transgenic fish expressed exogenous capsaicin receptor under a Purkinje cell-specific promoter, alongside a pan-neuronal nuclear-localized calcium indicator to validate activation specificity and register anatomy (**Fig. 1B**; *Tg(aldoca:TRPV1-tagRFP; elavl3:H2B-GCaMP7f; nacre−/−)*)^26,27^. Because zebrafish possess no endogenous capsaicin receptor, systemic application resulted in highly specific, inducible activation of Purkinje cells (**Fig. 1C**).

**Figure 1.**
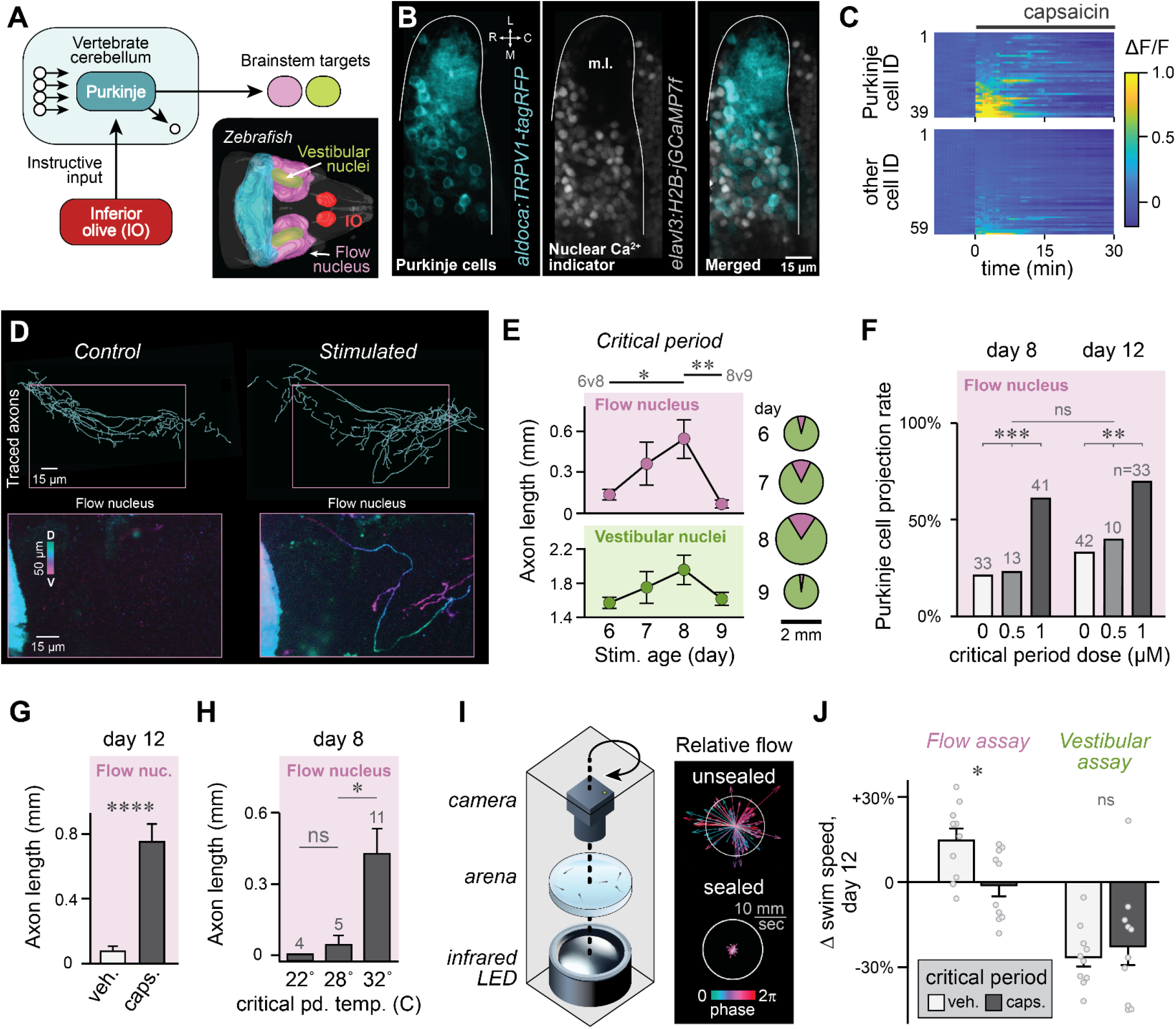
Activity-dependent plasticity of Purkinje cell projections in zebrafish. (**A**) Schematic of the vertebrate cerebellar learning circuit and rendering of zebrafish circuit anatomy. (**B**) Zebrafish cerebellum (*outlined*; 12 days) showing cytosolic TRPV1 expression (*cyan*) in Purkinje cell somata and dendrites in the molecular layer (ml), as well as pan-neuronal nuclear jGCaMP7f expression (*gray*). (**C**) Calcium dynamics of cerebellar neurons grouped by TRPV1 expression during application of capsaicin (6 days, *Tg(aldoca:TRPV1-tagRFP; elavl3:H2B-jGCaMP7f*)). (**D**) Horizontal projections of traced Purkinje cell axons in representative genetic control (*left*) and capsaicin-exposed (*right*) brains. Depth-coded projections (*bottom*) show axons throughout the flow nucleus following stimulation (12 days). (**E**) Length of Purkinje cell axons in flow nucleus (*top left*) and vestibular nuclei (*bottom left*) at 12 days following 7-hour capsaicin exposure at 6, 7, 8, or 9 days (*n*=9-10). Pie charts reflect relative length of projections to flow (purple) and vestibular nuclei (green) and are scaled by total length of projections. ^*^ *p*<0.05, ^**^ *p*<0.01 by Tukey posttest after two-way ANOVA. (**F**) Proportion of brains with flow nucleus innervation by Purkinje cells as a function of capsaicin dose, measured at end of stimulation (8 days, effect of dose, ^***^ *p*<0.001) or at 12 days (effect of dose, ^**^ *p*<0.01). Group sizes above bars. (**G**) Length of Purkinje cell axons (mean ± SEM) in the flow nucleus (*F*, ^*\*\*\*\**^ *p*<0.0001) in stimulated and control fish (*n*=8; 12 days). (**H**) Length of Purkinje cell axons in the flow nucleus as a function of temperature during capsaicin stimulation (8 days). (**I**) Schematic of behavioral assay with arena orbiting in register with infrared camera and backlight (*left*). Freely-swimming larvae were recorded in unsealed or sealed conditions, with attendant flow dynamics quantified using particle image velocimetry (*right*). Vectors convey flow horizontal velocity by orbit phase. (**J**) Effect of prior Purkinje cell stimulation on change in swimming speed from baseline due to orbital motion in unsealed (*left*, flow assay) or sealed conditions (*right*, vestibular assay; 2 runs for each of 5 groups of 8 fish plotted by run, with overall mean).

Stimulation during a brief critical period caused lasting effects on long-range projections of Purkinje cells. We stimulated Purkinje cells with capsaicin for 7 hours on 6, 7, 8, or 9 days (post-fertilization) and then examined morphology at 12 days. Purkinje cell axons were highly sensitive to stimulation during a specific developmental window (**Fig. 1D**). Following stimulation at 7 or 8 days, Purkinje cell axons grew within primary sensory areas for head acceleration, the vestibular nuclei, as well as for water flow, the medial octavolateral nucleus (‘flow nucleus’; **Fig. 1E**). Stimulation caused significantly more growth at 8 than at 6 or 9 days (n=9-10; two-way ANOVA, main effect of stimulation age, F3,67=5.00, p<0.01; Tukey posthoc tests of pairwise ages, 6v8 days: *p*<0.05, 8v9 days: *p*<0.01, all else *ns*). Given low baseline innervation of the flow nucleus, stimulation-induced growth biased Purkinje cell projections there (mean fraction in flow nucleus: 7.6% after stimulation at 6, 14.5% at 7, 19.6% at 8, and 3.8% at 9 days). We conclude Purkinje cell projections are regulated by activity for a brief window early in the life history of a zebrafish.

To augment plasticity, we stimulated Purkinje cells chronically for 48 hours throughout the critical period. By the end of stimulation at 8 days and persisting until 12 days, stimulation dose-dependently increased the proportion of brains with Purkinje cell projections to the flow nucleus, from approximately 1/4 to 2/3 at a dose of 1 µM capsaicin (**Fig. 1F**, significant effect of dose (χ^2^(1)=22.62, *p* < 0.001) but not age (χ^2^(1)=2.73, *ns*). Furthermore, by day 12, critical period stimulation increased the length of Purkinje cell axons within the flow nucleus by an order of magnitude (**Fig. 1G**, 76.0 vs. 753.7 μm; n=8, t-test, t_14_=5.97, *p*<0.0001). To confirm capsaicin signaled via exogenous capsaicin receptor, we adjusted incubation temperature, which co-modulates channel conductance^28,29^, and found that warming could augment the effect of stimulation at day 8 (**Fig. 1H**, *n*=4-11; one-way ANOVA, effect of temp.: F_2,17_=5.52, *p*<0.05; Bonferroni: 28 vs 32C, *p*<0.05; 28 vs. 25C, *ns*). Elongated axons appeared immature with small, sparse boutons (**Supp. Fig. S1**) and even sporadically exhibited ectopic projections to the spinal cord, caudal hindbrain, and contralateral cerebellum (**Supp. Fig. S2**). Finally, we observed thinning of the molecular layer that comprises Purkinje cell dendrites, consistent with results in other models (**Supp. Fig. S3**)^30,31^. Activity during the two-day critical period can therefore drastically remodel Purkinje cell anatomy, particularly their long-range projections to the brainstem.

Growth of Purkinje cell axons in brainstem sensory structures corresponded to altered sensory responses. Because swimming is suppressed by inhibition of neuropil within the flow nucleus^32^, where GABAergic Purkinje cell axons heavily projected after stimulation, we designed a behavioral assay to compare swimming activity in response to flow and vestibular stimuli. While filming swimming behavior for posthoc image processing with machine learning, we applied orbital motion to achieve chronic variable acceleration and reversibly suppressed water flow by damping surface waves, mitigating subsurface flow by an order of magnitude (**Fig 1I;** see *Methods*). Following critical period Purkinje cell stimulation, fish were hyposensitive to water flow. Although motion with strong water flow reliably increased swimming speed in control fish at 12 days (+14.6%), it failed to alter swimming following Purkinje cell stimulation (−1.1%; **Fig. 1J**; *n*=5 groups of 8 fish with 2 repetitions each, linear mixed effects model, F1,17=5.40, *p*<0.05). In contrast, Purkinje cell stimulation had little effect on responses to primarily vestibular stimuli (motion with damped waves), which consistently suppressed swimming (−26.5% in controls vs. −22.7% after Purkinje cell stimulation, *ns*). Therefore, Purkinje cell stimulation not only caused an order of magnitude increase in projections to the flow nucleus but also suppressed behavioral responses to water flow.

### The IO regulates experience-dependent plasticity of cerebellar projections

Since cerebellar projections are labile during a critical period, and IO instructive inputs rely on those projections to transform plasticity into learning, we next examined the role of the IO in the critical period. Olivary neurons are key regulators of Purkinje cell activity and provide potent depolarization across vertebrates^33,34^. Direct chemogenetic stimulation of Purkinje cells supersedes olivary drive, so we instead tested whether IO modulates plasticity of Purkinje cell projections due to experience. Both the vestibular and flow-sensing systems are activated by motion, and we previously noted adaptation to early motion experience in zebrafish^35^, so we placed fish under chronic motion (CM) during the critical period (**Fig. 2A**).

**Figure 2.**
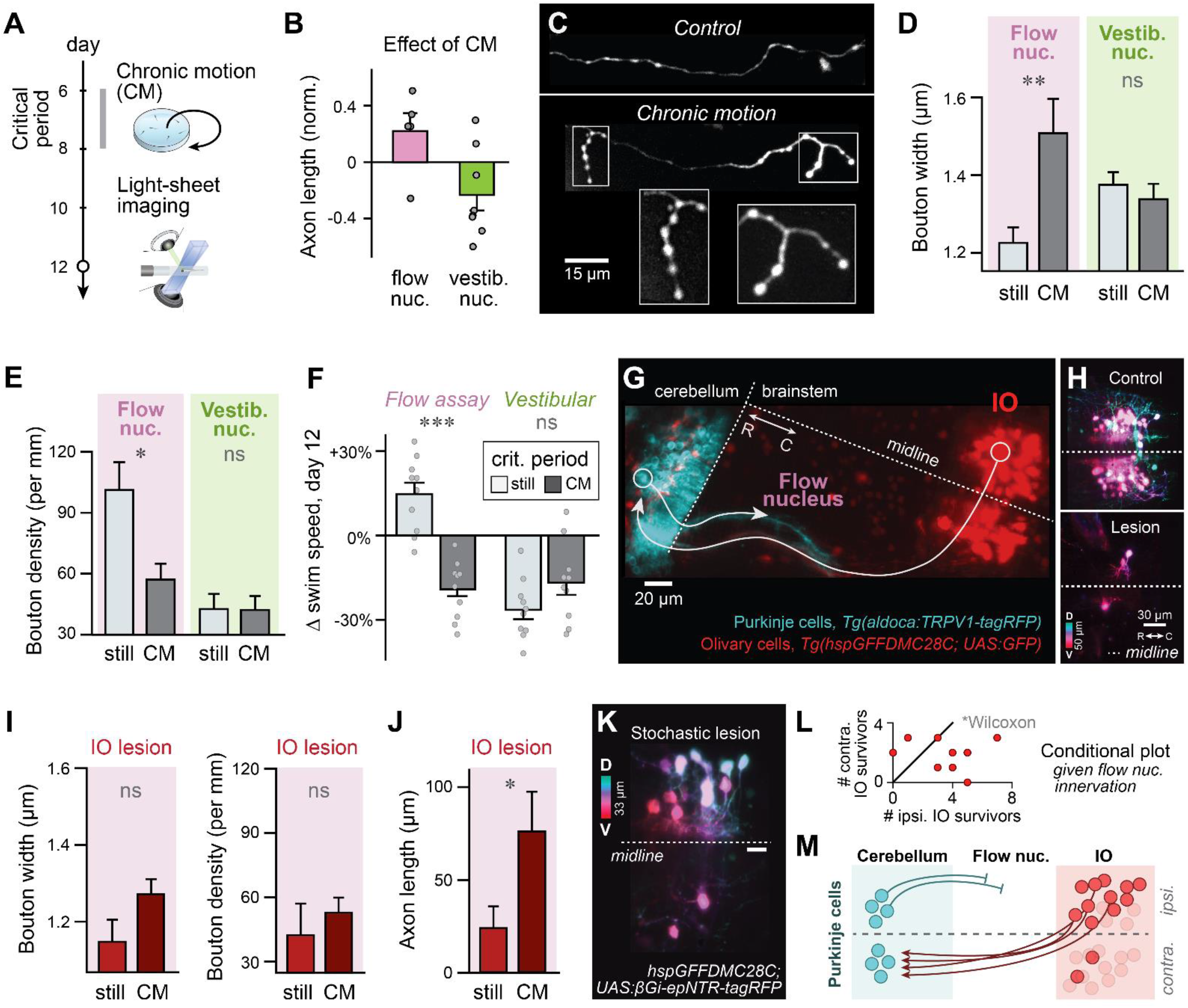
The IO regulates experience-dependent plasticity of Purkinje cell projections. (**A**) Fish were exposed to CM from 6-8 days before evaluation at 12 days. (**B**) Change in axon length (mean ± SEM) at 12 days in the flow nucleus (*pink*) and vestibular nuclei (*green*) from CM. *ns* by t-test. (**C**) Representative maximum intensity projections of Purkinje cell axons in the flow nucleus at 12 days with or without CM from 6-8 days. (**D, E**) Median bouton width (*D*) and density (*E*) on Purkinje cell axons in the flow nucleus (*left*) and vestibular nuclei (*right*) with or without CM (mean ± SEM). (**F**) Effect of CM from 6-8 days on swimming speed change at 12 days in response to orbital motion in unsealed (*left*) or sealed conditions (*right*; 2 runs for each of 5 groups of 8 fish plotted by run, with overall mean; ^***^ *p*<0.001). (**G**) Horizontal maximum intensity projection of transgenic for lesioning IO prior to CM and Purkinje cell tracing, here expressing a fluorescent reporter revealing climbing fiber apposition to Purkinje cell somata. (**H**) Representative, depth-coded projection of IO chemosensitizer expression (βGi-epNTR-tagRFP) without (*top*) and with (*bottom*) metronidazole-mediated cell ablation (12 days). (**I, J**) Axonal morphology at 12 days after IO lesion at 5 days and CM from 6-8 days. Median bouton width (*I, left*, ns by t-test), bouton density (*I, right*, ns), and total axon length (*J*, ^*^ p<0.05) are plotted as mean ± SEM. n=23-24 brains, of which 11 and 5 had axons for CM and controls, respectively. (**K**) Depth-coded representative micrograph of chance asymmetric IO lesion at 12 days. (**L,M**) For brains with at least one Purkinje cell axon innervating the flow nucleus, IO lesions were more thorough in the contralateral hemisphere, in terms of extant IO cells upstream of the flow nucleus projection (*L*, ^*^ p<0.05, Wilcoxon signed-rank test). Logic of conditional plot (*M*).

Flow nucleus projections and behavioral flow responses were sensitive to motion experience, providing a foundation to study olivary regulation. In contrast to direct stimulation with capsaicin, CM had only modest effect on the length of Purkinje cell axons within the flow and vestibular nuclei by 12 days (+22% in flow, −23% in vestibular nuclei; **Fig. 2B, Supp. Fig. S4**). However, axons specifically within the flow nucleus were strikingly different after CM (**Fig. 2C-E**). They exhibited larger boutons that appeared spatially clustered after CM (**Fig. 2C,D**; *n*=5-7; 1.51 vs 1.23 μm width for controls; t_10_=3.22, *p*<0.01) and were approximately half as dense (**Fig. 2E**; 57.1 vs 101.2 boutons per mm axon for controls; t_10_=2.49, *p*<0.05), indicators of axon maturity consistent with preferential pruning of small boutons (**Supp. Fig. S5**)^36,37^. In contrast, axons within the vestibular nuclei exhibited no change to bouton morphology or density from CM (**Fig. 2D-2E**; *n*=8; width: 1.34 vs 1.38 μm for controls; t14=0.73, *ns*; density: 42.1 vs 42.6 boutons per mm axon for controls; t14=0.04, *ns*). As with Purkinje cell stimulation, CM affected not only flow nucleus anatomy but also behavioral responses to flow (**Fig. 2F**). Specifically, CM prevented swim speed changes in the flow assay at 12 days (−19.0% vs. +14.6% for controls; *n*=5 groups of 8 fish with 2 repetitions each; Linear mixed effects model, F_1,17_=38.57, *p*<0.001), but had no significant effect in the vestibular assay (−16.6% vs. −26.5% for controls; F_1,17_=3.42, *ns*). We conclude early motion experience can drive maturation of Purkinje cell projections to the flow nucleus as well as behavioral flow sensitivity.

Experience-dependent plasticity of Purkinje cell projections was regulated by the IO. To test the necessity of IO for plasticity, we induced lesions of IO cells with a transgenic approach one day prior to CM. Specifically, fish expressed a chemosensitizer, nitroreductase, under an IO-specific promoter^38^, as well as fluorescently labeled Purkinje cells (**Fig. 2G,H**, *Tg(hspGFFDMC28C+; UAS:βGi-epNTR-tagRFP+; aldoca:TRPV1-tagRFP+; nacre−/−)*). Notably, IO lesions prevented effects of CM, preventing the increase in bouton width and decrease in bouton density in the flow nucleus (**Fig. 2I**; *n*=5-11; width: 1.27 vs 1.15 μm for controls; t14=1.68, *ns*; density: 52.7 vs 42.1 boutons per mm axon for controls; t14=0.74, *ns*; **Supp. Fig. S5**). Furthermore, although CM failed to alter axon length in the intact system, IO lesions revealed a capacity of CM to induce axon growth, causing a threefold increase (**Fig. 2J**, *n*=23-24, 76.0 μm CM vs. 23.9 μm for controls; t45=2.11, *p*<0.05). Effects of lesion were not systemic but pathway-specific, as Purkinje cells within individual brains were more likely to project to the flow nucleus if their candidate IO afferents, categorized by hemisphere assuming decussation, were more thoroughly lesioned (**Fig. 2K,L**; *n*=9, Wilcoxon signed-rank test, *p*<0.05). These results reveal a novel developmental function for the instructive IO, which typically orchestrates mature learning, to indirectly regulate wiring changes downstream of its cerebellar targets (**Fig. 2M**).

### Early experience activates IO and targets to direct Purkinje cell projections

To understand why CM drives maturation of Purkinje cell projections specifically to the flow nucleus and how the IO regulates this process, we recorded motion responses throughout the brain using calcium imaging. Fish sense motion with multiple sensory modalities that can activate the olivocerebellar system and its target structures, but neural responses to coherent motion are rarely described as stimuli are often decomposed by modality. Specifically, isolated water flow is transduced by the neuromasts of the mechanosensory lateral line and encoded in the flow nucleus, head acceleration is transduced by the vestibular endorgans that activate both the vestibular nuclei and IO, and visual motion activates the IO and putatively the vestibular nuclei (**Fig. 3A**)^39–44^, but the interaction of these brain areas during multimodal motion sensing is unclear. To record neural responses across the brainstem while imposing naturalistic motion, which requires displacement of the fish to activate the vestibular system, we devised an approach to record calcium dynamics of cells moving precisely within the focal plane of a microscope. To achieve such precision, we mounted a voice coil actuator on a linear stage atop a kinematic mount to calibrate travel within the focal plane. This device, the μSled, is open-source and low-cost, enabling reproducible and programmable translation of samples for intravital imaging (**Fig. 3B,C, Supp. Fig. S6 & S7** and *Methods* for fabrication details and performance validation). By moving fish with freed tails under a multiphoton microscope, we could apply coherent sensation of motion in vestibular, visual, and flow modalities, evidenced by deflection of a fluorescent neuromast (**Fig. 3D**).

**Figure 3.**
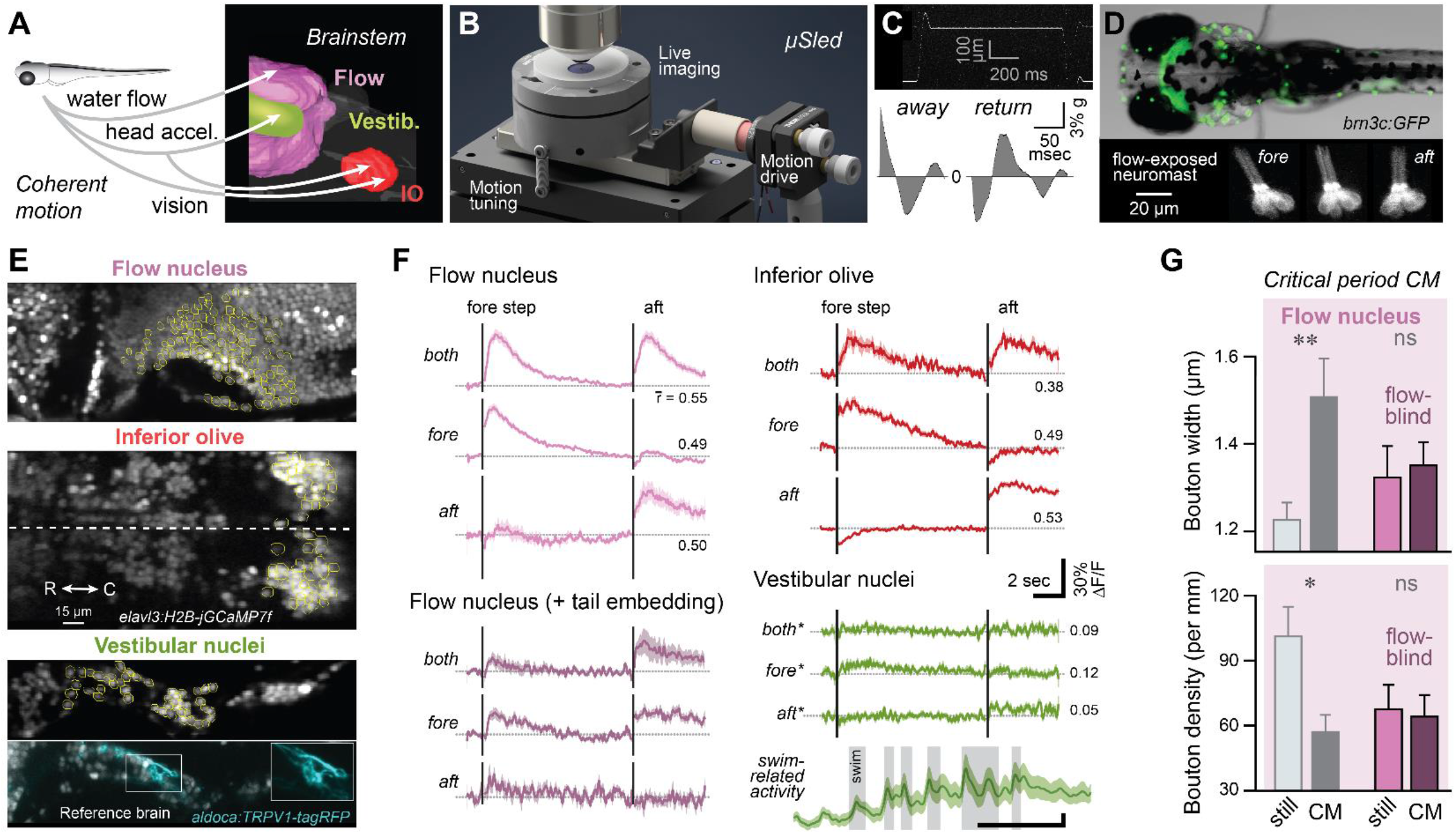
Neural responses to motion implicate coactivation with IO in target-specific maturation. (**A**) Schematic of known zebrafish brainstem encoding of modalities comprising natural motion. (**B**) Rendering of μSled, a device for tunable translation of samples to optically record neuronal motion responses. (**C**) Translation of a fluorescent microbead using μSled on a multiphoton microscope (*top*) used to calculate linear acceleration (*bottom*). (**D**) Fluorescent mechanosensory neuromasts (*Tg(brn3c:GFP)*) in a merged fluorescent and transmitted light micrograph (*top*) and neuromast deflection by the μSled. (**E**) Pan-neuronal, nuclear-localized calcium indicator (*gray*) with brain region-specific regions of interest (*yellow*) used to measure motion responses. Comparable vestibular nuclear anatomy is also presented in a second brain in which Purkinje cell axons are additionally labeled (*cyan*). (**F**) Calcium dynamics during μSled translation both fore (towards head) and aft (towards tail) within the flow nucleus, IO, and vestibular nuclei as mean ± SEM. Dynamics are shown for cells uniquely sorted into response type categories by correlation with stimulus-specific predictors (mean correlation within categories shown), and for the same flow nucleus responders before and after tail embedding to prevent water flow. ^*^Since no vestibular nuclear cells exhibited sufficient responses, dynamics are depicted for the 8 highest correlated cells within each category, as well as during periods of spontaneous swimming (*bottom right*). (**G**) Effect of ablating the mechanosensory lateral line on critical period CM-mediated axonal plasticity. Bouton width (*top*) and density in the flow nucleus (*bottom*) exhibited significant effects of CM (see Fig. 2D,2E) unless CM was applied in flow-blind fish after ablation of neuromasts (*ns*, t-test with Bonferroni correction, mean ± SEM).

Surprisingly, coherent motion strongly activated the IO and flow nucleus but not the vestibular nuclei. We imaged motion responses at 6 days in naïve fish and observed both directionally tuned and bidirectional neurons in both the flow nucleus and IO (see *Methods*). Of 265 flow nucleus cells imaged in one fish, 10.9% exhibited suprathreshold responses to both fore and aft motion, 20.4% responded to fore motion only, and 3.8% responded to aft motion only, and responsive cells in each category exhibited a mean correlation with stimulus predictors exceeding 0.49 (**Fig. 3E,F**). Similarly, of 149 IO cells, 4.0% responded to both fore and aft motion, 12.8% responded to fore motion only, and 40.3% responded to aft motion only, with mean correlation with stimulus predictors exceeding 0.38. In contrast, unlike in prior experiments of unimodal, oscillating head acceleration^45,46^, none of the 46 vestibular nuclear cells recorded in the Purkinje cell terminal fields exhibited any suprathreshold response to fore or aft stepwise motion (**Fig. 3E** *bottom* and **3F**). However, vestibular nuclear cells did activate strongly coincident with swimming behavior (**Fig. 3F**, *bottom*). These data suggest a potential explanation for IO-dependent maturation of Purkinje cell axons within the flow nucleus but not vestibular nuclei following motion experience (**Fig. 2D,E,I**): coactivation of flow nucleus and IO cells may promote maturation of intermediary Purkinje cell projections. To test this hypothesis, we decoupled flow experience from other modalities conveying motion. As expected, suppressing flow sensation by embedding the tail after imaging mitigated responses within the flow nucleus across each category of motion-responsive cells (**Fig. 3F**).

By interfering with flow sensation during CM, we identified coactivation of instructive inputs and targets as a potential mechanism for plasticity. Before exposing fish to CM during the critical period, at 5 days we lesioned lateral line neuromasts, creating ‘flow-blind’ fish throughout the critical period^35^. Without a functional lateral line to activate the flow nucleus, CM failed to increase bouton width or decrease density of Purkinje cell axons within the flow nucleus (**Fig. 3G;** *n*=9-10; width: 1.35 vs 1.32 μm for controls; t_17_=0.31, *ns*; density: 64.0 vs 67.5 boutons per mm axon for controls; t_17_=0.23, *ns*). Both axonal features in the flow nucleus of flow-blind fish were comparable to those of axons in the vestibular nuclei, which was not activated by motion and where CM fails to induce plasticity (compare to **Fig. 2D,E**). Furthermore, CM of flow-blind fish caused a tendency to grow axons within the flow nucleus, similar to CM in fish after IO lesion as well as direct Purkinje cell stimulation (*n*=19-24; 79.2 vs 34.4 μm for controls; t_41_=1.69, *ns*; data not shown). These data suggest experience-dependent maturation of Purkinje cell projections depends not only on IO activation but also on activity within target structures.

### Developmental plasticity can promote credit assignment

The discovery that Purkinje cell projections are highly plastic reveals a potential strategy not for solving credit assignment directly but for building circuits in which local plasticity can assign credit. Unlike artificial networks trained by backpropagation, which can compute each synaptic change from the entire downstream circuit, biological synapses are modified by local rules using signals available at the synapse and so depend on downstream architecture to make those local changes beneficial^2^. Accordingly, an instructive input to the cerebellum is thought to improve behavior because it specifically acts on Purkinje cells whose downstream function corresponds to that instruction’s tuning, an example of a widespread circuit motif for enabling credit assignment^3–5,47^. We propose an instructive input can construct the very circuit in which it later performs credit assignment, shaping downstream connectivity indirectly so local plasticity during learning can improve behavior. Specifically, our experiments suggest the IO could serve this dual purpose, as Purkinje cell axons matured within the flow nucleus only when both the upstream IO and downstream flow nucleus were activated by experience (see **Figs. 2M, 3F, 3G**). This dependence of Purkinje cell development on structures both up- and downstream is reminiscent of the correlation-based, Hebbian plasticity that organizes developing sensory systems^22^, suggesting the IO could build IO-Purkinje-target correspondence by an analogous rule.

To test whether developmental plasticity can construct circuits in which local plasticity suffices for credit assignment, we devised a computational model. Given that zebrafish use flow sensation and the cerebellum to control turning^48,49^, we tasked an olivocerebellar circuit with learning to move an agent about a plane by commanding two-dimensional effectors (**Fig. 4A**). In order to learn, IO cells with precise directional preference (matrix *A*_*io*_) conveyed feedback about performance to instruct heterosynaptic plasticity at Purkinje cell motor inputs (matrix *W*_*1*_), constituting a canonical Marr-Albus-Ito ‘learning rule’ ^50^. However, the circuit was initially pseudorandomly connected and first relied on another form of plasticity, a Hebbian ‘wiring rule,’ by which the IO strengthened Purkinje cell projections to coactive targets (matrix *W*_*2*_, see *Methods*). For the learning rule to improve behavior, that is for the circuit to perform credit assignment, IO error signals must be routed to Purkinje cells in which plasticity biases movement in the same direction. Whether the learning rule can improve behavior therefore depends on Purkinje cell projections established during development.

**Figure 4.**
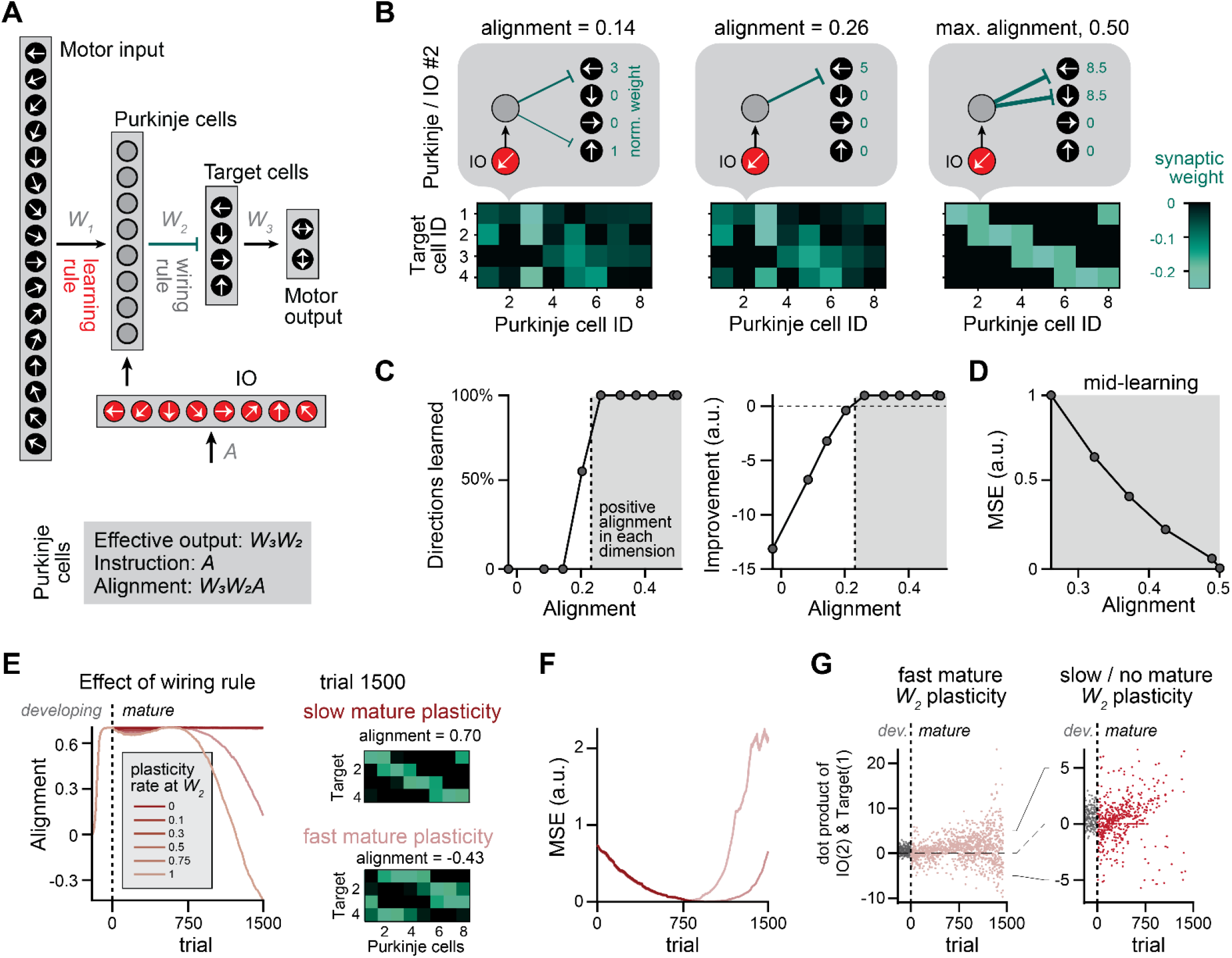
Cerebellar development can construct optimal architecture for learning. (**A**) Diagram of simulated cerebellar circuit. Purkinje cell inputs sparsely encode the direction of intended movement of an agent (see *Methods*). IO neurons convey the error as a function of direction, with each IO neuron projecting to one Purkinje cell. Arrows in cells reflect either intended movement direction (in input layer), error tuning matrix *A*_*io*_ (in IO), or motor function (in target cells and output layer). (**B**) Example connectivity patterns between Purkinje and target cells (*W*_*2*_) with schematic of a single Purkinje cell (*top*) and matrices of full Purkinje cell layer (*bottom*). Color represents synaptic weights between inhibitory Purkinje and target cells. (**C**) Effect of alignment from randomizing *W*_*2*_ on performance. For circuits with large positive alignment, learning converges such that all instructed directions are learned (*left*). At small positive or negative alignment, learning diverges such that the error increases with training (*right*). The vertical dashed line denotes the alignment in these arbitrary circuits at which both eigenvalues become positive. Because we impose *W*_*1*_ to be positive only, the circuit requires greater alignment to learn all directions (see **Supp. Fig. S8**). (**D**) Learning is faster with greater alignment, resulting in faster reduction of the mean squared error (MSE). (**E,F**) The wiring rule acting at *W*_*2*_ initially drives the circuit to maximum alignment but eventually reduces alignment after learning at *W*_*1*_ begins at trial 0 (*E*). Performance tracks alignment over the course of training, improving as the initially aligned circuit trains but then degrading as alignment is disrupted (*F*). (**G**) Correlation of activity between an IO and target cell that share the same directional tuning, measured through the dot product. During development (*gray*) this pair is strongly positively correlated, but not after the learning rule activates and the circuit becomes driven by motor input (*red*, trials > 0). If the wiring rule remains highly plastic while the learning rule is active, once correlation of IO and target cell activity cease to reflect alignment, learning fails (*left*).

For the learning rule to reliably improve performance, the connectivity must satisfy a constraint we derived analytically (**Appendix**). Although cerebellar anatomy exhibits a common motif enabling credit assignment, in which individual Purkinje cells bridge approximately matched instructions and downstream function, how connectivity impacts learning at population scale remains unknown. We show mathematically that for a cerebellar circuit to learn stably, there must tend to be directional similarity, or alignment, between IO tuning (*A*_*io*_) and the effective output of Purkinje cells mediated by the downstream circuit; with this circuit architecture, effective output is captured by the product *W3W_2_*, but results generalize to any arbitrary downstream architecture. Specifically, the instructive input and effective output of the Purkinje cell population must be positively aligned along every dimension; formally, the matrix *B*=*W*_*3*_*W*_*2*_*A*_*io*_ must be positive definite, meaning all the eigenvalues of the symmetrized matrix, *B+B*^*T*^, are positive (**Appendix, conclusions 1 & 3**). Thus, given a fixed wiring of a Purkinje cell to instructive inputs (*A*_*io*_) and fixed function of candidate targets mediated by their own downstream connectivity (here, *W*_*3*_), any Purkinje cell wiring to targets (*W*_*2*_) that satisfies the alignment condition ensures credit assignment. In other words, Purkinje cells here must influence movements in directions represented by their instructive inputs in aggregate, but each Purkinje cell need not exhibit positive alignment for the circuit to learn. Accordingly, a perfect alignment solution 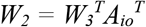 could result from Hebbian plasticity (**Appendix, Part B**). These findings suggest a development rule would promote local credit assignment if it strengthens connections from Purkinje cells to targets contingent on coactivation of targets and IO inputs of the Purkinje cell. The theory was confirmed in simulation of circuits with random architecture, as only positively aligned circuits learned the task (**Fig. 4B,C; Supp. Fig. S8A,B**) and the more aligned the circuit, the faster it learned (**Fig. 4D; Supp. Fig. S8C**). Therefore, simply by biasing the circuit toward positive alignment, developmental plasticity would tend to make future learning faster and the learning rule more viable.

Finally, we asked how developmental plasticity and learning interact. We found that development of Purkinje cell projections can promote credit assignment, but only if the wiring rule is restricted to a critical period and slowed below the rate of learning. In these simulations, the wiring rule promoted positive alignment because the IO signaled feedback about movements generated by spontaneously active Purkinje cell targets, affording the requisite activity correlation (see *Methods*), which is a distinct mechanism to achieve coactivation of IO and Purkinje cell targets from CM in experiments (**Fig. 3F**). Initially, the wiring rule drove the circuit towards maximal alignment (**Fig. 4E**, trial 0), enabling performance improvement via the learning rule simply by pre-configuring Purkinje cell projections (**Fig. 4F**, trial <750; **Supp. Fig. S8D-G**). Once learning began, however, the wiring rule became destructive.

Specifically, in successfully aligned circuits the learning rule reduced the component of error predictable from Purkinje cell motor inputs, so the IO signal became decorrelated from other Purkinje cell inputs (**Fig. 4G**)^51^. Therefore, if the wiring rule persists after successful learning, the resulting loss of correlation between Purkinje cell target activity and IO activity degrades alignment (**Fig. 4E,F**). By contrast, when the two sites of plasticity were appropriately balanced, with the wiring rule operating at less than or equal to half the rate of the learning rule, this degradation became self-correcting; as performance fell, the return of predictable error restored the correlation between IO and Purkinje cell targets, and the slow wiring rule acted on that restored correlation to gradually recover alignment. Thus, a key benefit of slowing plasticity upon critical period closure is to preserve the projection patterns established during development as learning proceeds. Collectively, our experiments and computational analysis suggest instructive inputs can use developmental plasticity mechanisms to construct and adapt essential architecture for credit assignment.

## Discussion

Our results outline a developmental solution to credit assignment in the cerebellum. In canonical cerebellar learning, instructive inputs to Purkinje cells from the IO heterosynaptically modulate the strength of other inputs^52–24^. However, these modifications are thought to improve behavior only if the downstream influence of a Purkinje cell is aligned with the instruction conveyed to it by the IO^7,10,11^. How the cerebellum establishes this alignment between the plasticity and output of its cells has been a mystery^55^. Our experimental and modeling results reveal that the developing cerebellum can achieve architecture for local credit assignment when developmental plasticity, orchestrated by experience-dependent activity, modifies Purkinje cell efferent connectivity such that downstream function becomes aligned with instruction. This alignment is required of the population but not single cells, such that a misaligned Purkinje cell that modulates synaptic weights counterproductively during learning is outweighed by the aligned majority. Notably, this biological mechanism approximates backpropagation of errors from machine learning while inverting its logic. Backpropagation computes synaptic changes tailored to an arbitrary downstream network^56,57^; here, error is assigned without knowledge of the downstream network by IO inputs of fixed tuning and routing, and instead the loss landscape becomes shaped around those instructions during development. In other words, by modifying downstream connectivity during development, instructive inputs can alter the function of target cells to accommodate learning through a simple, local plasticity rule.

Cerebellar developmental plasticity parallels activity-dependent wiring in sensory systems, where patterned activity can construct labelled lines across layers and regions^22,58,59^. In contrast to sensory system development, though, experience-dependent plasticity of the cerebellum was regulated by a dedicated instructive input. Purkinje cell connectivity is known to be affected by activity and developmental experience^24,30,31,60^, but less established is how these processes regulate the registration across Purkinje cells of IO instruction and target function. Here, Purkinje cell activation drove broad innervation of less saturated targets, exuberant growth that may provide a substrate for selective pruning^61^. The anatomical specialization of IO innervation, the climbing fiber, induces pronounced depolarization that drives Purkinje cell plasticity, even in developing zebrafish^34^, which may afford robust signaling for feedforward plasticity. The early arrival of developing climbing fibers relative to other Purkinje cell inputs^62,63^ could afford a privileged window to shape efferent connectivity in the nascent circuit. A winner-takes-all competition which prunes less active climbing fibers may complementarily preserve the input best suited to regulate downstream wiring and thereby promote credit assignment^60^. Consistently, climbing fiber responses are present within the first days of Purkinje cell differentiation in zebrafish, climbing fiber pruning accompanies the tuning refinement we observe across the critical period, and the IO is required for the emergence of functional cerebellar output in developing mammals^64–66^. Critical period closure may mark a transition in the plasticity the IO most prominently induces, from a Hebbian-like rule that strengthens connections between co-active partners across developing brain regions to the mature, anti-Hebbian plasticity that supports cerebellar learning.

Confining plasticity to a brief critical period carries profound benefit. Simulations revealed a computational cost of leaving the critical period open by maintaining highly plastic projections of Purkinje cells amidst learning. If developmental plasticity continued at high rates, it corrupted the very architecture it previously established that made learning possible, so closing the critical period preserves downstream architecture and renders learning robust. In turn, it becomes clear why the plastic window opens when it does in the context of the animal’s ecology and life history. The critical period falls during the first few days of active hunting, when larval zebrafish, like other teleosts entering external feeding, face intense mortality risk from predation and starvation^67–70^. Because behavior at this stage bears so directly on feeding success and predator evasion, the short-term benefits of rapid, persistent remodeling likely outweigh any long-term costs^71,72^. Consistently, the zebrafish cerebellum assembles rapidly and regulates learning and locomotion within one week of fertilization, and olivary sensory tuning emerges early, supplying the physiological preconditions for the plasticity we describe^48,64,73–77^. The window therefore opens once the basic circuit is assembled but while behavioral consequences for fitness remain high, then closes before persistent remodeling might degrade the acquired architecture. Like the experience-dependent plasticity that tunes sensory circuits to the statistics of the sensed world, including when those statistics are atypical^78,79^, this mechanism adapts the cerebellum and flow processing to the animal’s early motion experience. Purkinje cell projections to the flow nucleus have been described only rarely^80^, which may reflect the impoverished experience of typical laboratory rearing in stagnant water^35^.

To establish suitable connectivity for credit assignment, the mechanism investigated relies on activity correlations that reflect the causal relationship between action and sensory feedback. The sensorimotor loop here was closed by olivary encoding of reafference, the sensory feedback an animal generates through its motor output. Through reafference, active targets of Purkinje cells became correlated with olivary cells whose sensory tuning was aligned with the targets’ ultimate motor consequence. In other words, developmental plasticity facilitated subsequent motor learning in simulations because the loop conveyed to Purkinje cells how downstream connections function in sensory space. In this way, a solution to credit assignment arises from registration of sensory feedback with motor function, reflecting causality at the level of individual hidden layer neurons. In experiments, where the sensory experience was imposed and not reafferent, activity also drove circuit remodeling. Motion activated both the IO and flow nucleus, and flow sensory blockade mitigated flow nucleus responses as well as plasticity. A parsimonious interpretation is that CM co-opted a typical developmental process, artificially coactivating the IO and flow nucleus targets to bias projections. If IO activation instead conveys reafference such that correlations across regions become grounded in the physics of the body and the organization of sensory and motor maps, a circuit could achieve credit assignment while accommodating individually variable motor plants or sensorimotor maps. Critically, no purely internal scheme can originate this registration; climbing-fiber collaterals to the cerebellar nuclei, for example, are strongest during development and likely regulate circuit formation^81,82^, but they could only serve to register Purkinje efferents with function-matched targets if the olivonuclear projection was itself already registered, relocating rather than resolving the problem.

What local plasticity rule could mediate the empirical effects of early-life experience and simulated developmental wiring? We propose a Hebbian-like, correlation-based rule of the kind that drives activity-dependent wiring throughout the developing brain^22,59,83^, operating here to link neurons of common directionality in task space, both in terms of IO sensory tuning and efferent neuron function. Such co-tuning is well documented in the mature cerebellum, including in oculomotor regions where IO cells and Purkinje cell targets share directionality during smooth pursuit and saccade^84–86^. Given that prominent co-tuned cerebellar organization emerges during development in our simulations, we speculate this plasticity rule may not only afford adaptation to experience but also construct canonical cerebellar architecture; operating solely on naturally tuned responses during typical mammalian development, this rule could build the aligned IO–efferent organization of circuits such as the oculomotor vermis. Organizing projections by aligned motor function in this way also grouped cerebellar output into synergies (**Fig. 4B**), a developmental route for solving a central problem of motor coordination^87^. This developmental plasticity rule may bridge reafferent delays so IO activity can promote appropriate connectivity downstream, as the cerebellum can precisely regulate plasticity using long-latency feedback^88,89^ and immature circuits tend to employ broad integration windows driving potentiation with negative lags^90–93^. Consistent with broad temporal integration for early wiring, immature IO neurons low-pass filter excitatory synaptic input^94^.

Beyond realizing feedback-guided wiring in a biological circuit, obviating backpropagation’s implausible weight transport^2,95^, this developmental strategy affords economy of wiring. Our development mechanism and feedback alignment, the algorithm first proposed to circumvent weight transport using feedback, both serve to align weight updates with the true loss gradient. However, feedback alignment uses distinct feedback pathways to train multiple layers simultaneously, whereas this developmental mechanism reuses a single feedback pathway to train both layers sequentially. Plasticity at both sites for development and learning need not be sequenced, as the critical period could remain open without stifling learning if developmental plasticity was slow relative to the canonical cerebellar learning mechanism. Regardless, preconfiguring downstream connectivity during development restricted learning-related plasticity to a single layer, which accelerated convergence in our simulations consistent with networks trained using feedback alignment^96^.

Our formalization implements developmental plasticity specifically in the projections of Purkinje cells, which adapt to fixed wiring of IO input and connectivity further downstream. Mathematically, the alignment condition could equally be met by adjusting IO input or Purkinje target function, but only the Purkinje cell efferent synapses have access to both IO and target cell activity, so a local, Hebbian-like rule can assemble the network only there. Furthermore, the octavolateral and vestibular targets of Purkinje cells are evolutionarily ancient brainstem structures closely tied to the origin of the cerebellum^14,25^, so cerebellar projections plausibly evolved and may develop to engage established downstream circuits rather than the reverse.

The formalization also relies on deliberately simple plasticity rules, chosen to prove sufficiency of learning. While the nature of the signal encoded by the mature climbing fiber is still intensively studied^97–100^, here we adopted the canonical error-based cerebellar rule for its convergence and analytical tractability, as success is readily quantified. Our simulations also assumed linear Purkinje cell responses for simplicity, consistent with their high basal firing rates, and we prove that for linear networks a positive alignment between instructive input and Purkinje cell effective output is sufficient for successful learning. The result extends to nonlinear networks (**Appendix, conclusion 2**), but alignment for these networks must be exact because the network’s greater flexibility necessitates stronger constraint on the pre-trained downstream layer.

The credit assignment problem arises for any deep network in which local synaptic changes must serve global performance, and it remains the central obstacle to understanding learning in biological networks^2^. Error backpropagation, sufficient for artificial systems, cannot be implemented by the brain, and the schemes proposed to solve credit assignment biologically establish plausibility without an empirically identified substrate. Here we identify such a mechanism in vertebrate development, in which instructive inputs reinforce suitable downstream architecture for subsequent credit assignment. Learning with dedicated instructive inputs is not peculiar to the cerebellum, and developmental regulation of downstream connectivity could plausibly foster alignment across learning mechanisms and circuits, potentially including dendritic plateau potential-instructed plasticity in hippocampus and neocortex^101–103^. A single developmental mechanism could afford representational flexibility while also preconfiguring networks for credit assignment through precise routing of instructive inputs among populations of heterogeneous function. Together, these results reinforce the notion that training neural networks, both biological and artificial, requires mechanisms to not only guide local plasticity but also instantiate architecture for its viable expression^104^.

## Methods

### Animals

#### Zebrafish husbandry

Zebrafish *(Danio rerio*) embryos were naturally spawned, collected, and maintained in standard E3 medium (0.30 M NaCl, 0.01 M KCl, 0.03 M CaCl_2_, 0.02 M MgSO_4_, pH 7.0) refreshed daily to maintain water quality. Fish without inflated swim bladders by 4 days, or days post-fertilization (dpf), were excluded from all experiments. Embryos and larvae were housed at 28°C under a 14-hr light / 10-hr dark cycle at densities of 20-40 fish per 100-mm Petri dish. Fish were transferred to 150 mL of fresh E3 at 8 dpf and fed a commercially available powdered fry diet daily. All experiments were performed in accordance with protocols approved by the Institutional Animal Care and Use Committee (IACUC) of the College of Letters and Science at the University of Wisconsin-Madison.

#### Transgenic lines

All animals were homozygous for the *nacre* mutation, which impairs the *mitfa* gene required for melanophore formation, resulting in transparent skin that facilitates optical access to the brain^105^. For imaging Purkinje cells, unless otherwise indicated, experimental animals were homozygous for *Tg(aldoca:TRPV1-tagRFP)* and heterozygous for *Tg(elavl3:H2B-jGCaMP7f)*^106^. The *Tg(aldoca:TRPV1-tagRFP)* line expresses rat TRPV1 selectively in all cerebellar Purkinje cells, enabling targeted chemogenetic activation via bath application of capsaicin^27^. *Tg(elavl3:H2B-jGCaMP7f)* was chosen for pan-neuronal expression of genetically encoded calcium indicator with nuclear localization, which streamlines cell segmentation. Genotypes were confirmed by fluorescence at 5 dpf. For ION lesions, we used *Tg(hspGFFDMC28C)*, a Gal4 driver line with specificity for the inferior olive^38^. This line was crossed with *Tg(UAS:βGi-epNTR-tagRFP)* to express an enhanced-potency nitroreductase (epNTR), allowing inducible chemogenetic ablation of climbing fiber cerebellar inputs upon metronidazole exposure^107^. Lesion-study animals were offspring from an incross of *hspGFFDMC28C; UAS:βGi-epNTR-tagRFP; aldoca:TRPV1-tagRFP*.

### Chronic motion (CM)

Fish were visually screened at 5 dpf for expression of *Tg(aldoca:TRPV1-tagRFP)* and *Tg(elavl3:H2B-jGCaMP7f)*. Then at 6 dpf, they were randomly assigned to one of four conditions: (1) E3 medium + stagnant water, (2) E3 medium + CM, (3) capsaicin + stagnant water, or (4) capsaicin + CM. Capsaicin (1 μM; final DMSO <0.05%) was delivered by immersion in 25 mL of solution in 100-mm Petri dishes; vehicle-only controls were handled identically, and all solutions were refreshed after 24 hr. Mechanical stimulation was provided by a tilting shaker-incubator oscillating ±7° at 0.33 Hz (Incubator Shaker II, Boekel). In a subset of CM-exposed fish (6/19 in the 12 dpf study), an orbital shaker in an incubator was instead used (Labline 3520, 60 rpm, 3/4” circular path). Measures of axon length, bouton density, and bouton size did not differ between flow conditions (p > 0.15 for all comparisons) and results were pooled. All groups were maintained in the same incubator at ~28°C before and after 48 hr treatment (6-8 dpf), transferred to fresh E3 medium after CM and fed daily until imaging.

### Light Sheet Microscopy

Purkinje cell anatomy was visualized using a lab built Flamingo light sheet fluorescence microscope being developed and maintained by the Huisken and Eliceiri groups at the Morgridge Institute for Research equipped with a Nikon 16x/0.8 detection lens with a 400mm tube lens for 32x magnification. Fish were vertically mounted in 2% low-melting point agarose (Thermo Fisher Scientific 26520) within glass capillaries (1.0/1.5 mm) by retracting a plunger into the capillary. After agarose solidification, samples were dispensed into an E3-filled chamber and positioned just outside the capillary in the optical path for imaging^108^. Z stacks were acquired using a 561 nm laser to excite TRPV1-tagRFP and a 488 nm laser to excite nuclear-localized jGCaMP7f. A T-SPIM configuration was used for the microscope with two opposing illumination arms and a single orthogonal detection arm. Imaging was performed at 12 dpf unless otherwise specified, a stage at which fish maintain optical transparency while allowing assessment of prolonged treatment effects on Purkinje cell anatomy.

### Axon Tracing

#### Length quantification

Axonal lengths were manually traced and regions defined in Fiji by experimenters blinded to experimental group using Simple Neurite Tracer (SNT), an ImageJ plugin for semi-automated tracing of neuronal processes in three-dimensional image stacks^109,110^. Tracing was performed on z-stacks acquired with light sheet microscopy. Each axon was traced in three dimensions to its visible terminal point, and only clearly distinguishable axons were included. Flow nucleus (medial octavolateral nucleus) innervation lengths were quantified by summing traced axon segments within flow boundaries as defined in the mapzebrain atlas^111^; the region had a rostral boundary of the rostral extreme of the hindbrain and was caudally bounded by the white matter tract labeled “Neuropil Region 2” in the Z-Brain atlas^112^. The dorsal-ventral boundary extended from the dorsal brain surface to the ventral edge of the corpus cerebelli. Axon segments below this limit were excluded. Laterally, the region encompassed the lateral three-fourths of the brain at the dorsal extreme, tapering to the lateral one-third at the ventral extreme. Vestibular nuclei axon lengths were quantified by summing traced segments within the lateral and tangential vestibular nuclei, defined as lateral to Neuropil Region 3 (Z-Brain atlas) extending from the dorsal limit of the tangential vestibular nuclei to the ventral extreme of the spinal backfill vestibular population. The rostral boundary was set caudal to a point one Mauthner cell body width rostral to the Mauthner cell itself.

#### Bouton quantification

Putative boutons on Purkinje cell axons were first identified as bright swellings of the local axon. Swelling widths were measured in Fiji on image stacks of standardized brightness using the ROI Manager tool to draw a chord in the plane with widest swelling at the widest point orthogonal to the axon at that location^109^. To account for differences in axon labeling and width, after swelling measurements those exceeding a width threshold of one pixel greater than the mean axon diameter (949 nm) were classified as boutons. Then, boutons were counted only on axons previously marked in Simple Neurite Tracer (SNT) reconstructions to ensure correspondence with axonal length measurements. All tracings were performed blind to treatment.

### Behavioral Analysis

#### Behavioral assay

Behavioral assays were conducted on 12 dpf fish with or without CM from 6-8 dpf. Experiments were carried out on freely swimming zebrafish during the light phase of the 14:10 hr light/dark cycle. Behavior was recorded at 105 frames per second using a custom-built apparatus consisting of an orbital shaker (Labline 3520, 60 rpm, 19 mm circular path) and an overhead infrared digital camera (BFS-U3-162SM, FLIR) mounted on the moving platform to maintain a stable top-down view. Illumination was provided by a 5W, 940 nm infrared LED backlight (LED World) beneath the dish and an aspheric condenser lens to focus illumination (Thorlabs, ACL7560U-B, Ø75mm, F=60mm, NA=0.61), plus an overhead visible LED. Eight fish were recorded simultaneously in 60-mm diameter Petri dishes containing 42 mL of E3. Dishes were filled to maximum capacity with dish lids placed directly atop the water surface, sealing the dish without air bubbles. Following a 1-min acclimation, two 60-s recordings were obtained under static conditions, followed by a 1-min acclimation and two recordings with the shaker active (−flow). The lid was then removed to allow surface wave formation, and two additional recordings were obtained after a 1-min acclimation (+flow). This sequence was repeated for five rounds, alternating treatment groups within each round, with E3 replaced between rounds.

#### Behavioral tracking

Larval behavior was analyzed using Social LEAP Estimates Animal Poses (SLEAP), an open-source, deep-learning-based framework for multi-animal tracking^113^. A top-down network model was trained on 233 manually labeled frames with tracking focused on three anatomical landmarks: the caudal edge of the swim bladder and the rostral-most point of each eye. Behavioral metrics were extracted from tracked coordinates using custom MATLAB scripts, available on request, to low-pass filter paths to disentangle active swimming from shaker-frequency sloshing (**Supp. Fig. S9**).

#### Particle image velocimetry

To quantify water motion in the +flow and –flow behavioral conditions, the medium was seeded with tracer particles (d50 = 10 μm, specific gravity = 1.1; 110P8, Potter Industries) and illuminated with a continuous-wave laser diode expanded into a planar sheet using cylindrical optics (https://www.instructables.com/Low-Cost-PIV-System/). Image pairs were acquired on the orbital shaking behavioral rig and analyzed with PIVlab^114^. Spurious vectors were removed by applying a local spatiotemporal median filter with replacement.

### Lesion Experiments

#### ION lesion

Fish of genotype *hspGFFDMC28C; UAS:βGi-epNTR-tagRFP; aldoca:TRPV1-tagRFP* were screened at 5 dpf and treated with 10 mM Metronidazole for 24 hr^38,107^. Fish were washed in fresh E3 before experiencing CM from 6-8 dpf (or stagnation for genetic controls). Lesion efficacy was confirmed at 12 dpf using light sheet microscopy to examine ION somata, and Purkinje cell axonal lengths and boutons were quantified thereafter. Unlesioned controls were not traced due to additional fibers labeled by *Tg(hspGFFDMC28C)* that obscured Purkinje cell-specific tracing.

#### Lateral line ablation

Fish of genotype *aldoca:TRPV1-tagRFP; elavl3:H2B-jGCaMP7f* underwent lateral line ablation at 5 dpf by immersion in 50 μM copper sulfate for 60 min at room temperature, as previously^35^. After thorough washing in fresh E3, fish recovered for 24 h and then experienced CM from 6-8 dpf, then at 12 dpf imaging was performed to assess Purkinje cell anatomy.

### Calcium Imaging

#### Capsaicin response

Fish of genotype *aldoca:TRPV1-tagRFP; elavl3:H2B-jGCaMP7f* were imaged at 28.5°C during the light phase of the 14:10 h light/dark cycle using a Flamingo light sheet microscope. Baseline activity was recorded in E3 medium before exchanging the chamber medium for a 1 µM capsaicin solution in E3. Image sequences were acquired at 4.982 fps, with 64 frames collected at the start of each 45 s interval. Composite RFP/GCaMP images were generated to classify cells as RFP-positive (Purkinje cell) or RFP-negative (non-Purkinje cell) in the cerebellum. Time series were motion-corrected for lateral drift using the TurboReg plugin for ImageJ, applying a translation transformation based on a template maximum-intensity projection of stable frames^115^. Nuclear regions of interest (ROIs) were segmented using the EfficientViSAM-l2 implementation of the Segment Anything Model (SAM) plugin to obtain closely fitted cell boundaries^116^. Fluorescence traces were then extracted for each ROI, and *ΔF/F* was calculated by normalizing fluorescence changes to the baseline mean (*F*_*0*_) with custom code in MATLAB.

#### Motion responses with μSled

Measuring responses to naturalistic motion requires moving the fish in order to stimulate the vestibular system, previously accomplished by moving either specimen alone or an entire microscope^40,117^, alongside visual and lateral-line systems. We translated larvae within the imaging plane during calcium imaging using a custom, programmable sled (μSled, see **Supp. Fig. S6** for fabrication details). Translating the fish out of focus would change fluorescence for optical rather than physiological reasons, so we designed the μSled for tunable translation direction in order to travel precisely within the imaging plane. In-plane linear translation across the travel range was validated by imaging fluorescent microbeads (**Supp. Fig. S7**). The device uses a voice-coil actuator (VC125, Thorlabs) to drive a precision linear rail (BWU25-75; IKO, Japan) over a horizontal distance up to 10 mm but typically within a 200–500 µm range, with a light extension spring providing a centering force along the motion axis. A customized kinematic pitch/yaw mount (KPY1, Thorlabs) allowed fine adjustment of the orientation of the mounted larva, and an adjustable three-point mount (fine-thread M3 × 0.20 screws in threaded bushings, F3SS15/F3SSN1P, Thorlabs, contacting hardened seats held by stiff springs) maintained precise Z-axis positioning across the full travel range. The assembly was built into a two-part 3D-printed base (**Supp. Fig. S6**) permitting precise X–Z and Y–Z adjustment relative to the microscope, with voice-coil centering set by another kinematic mount (KMS, Thorlabs). Drive voltages were generated in the microscope’s Prairie View software and routed to a custom OPA541-based amplifier (Forward Scientific).

We imaged on a two-photon microscope (Bruker Ultima 2PPlus) in resonant-scanning mode, because two-photon excitation uses infrared light invisible to the fish, which allowed the visual environment to be controlled independently of imaging using an overhead red LED. Single planes were acquired sequentially, focused on the flow nucleus, vestibular nuclei, and inferior olive, identified anatomically using the pan-neuronal nuclear reporter (*elavl3:H2B-jGCaMP7f*, as for light-sheet imaging) with reference to the mapzebrain and Z-Brain atlases^111,112^. Lateral and tangential vestibular nuclei were bounded as in *Axon Tracing*. Larvae were mounted horizontally in agarose, with agarose surrounding the tail excised to form a 5 mm-wide channel while the head remained embedded, allowing water to flow over the tail during motion. Later, to mitigate visual or lateral line contributions to the motion response, the tail was re-embedded in agarose to suppress water flow or the LED was removed, respectively. Flow over the tail was confirmed by imaging bending of freed mechanosensory neuromasts of the lateral line in the *brn3c:GFP* line. For calcium imaging, fluorescence was measured at each stimulus position and ΔF/F computed against a position-specific baseline rather than a single global baseline to accommodate field nonuniformity. Motion correction, ROI segmentation, and ΔF/F calculation otherwise followed the calcium-imaging pipeline described for light sheet microscopy. Per-cell responses were averaged across stimulus repetitions (5–10 trials) following causal Butterworth filtering in MATLAB.

To identify motion-responsive cells, stimulus regressors were generated by representing each fore and aft stepwise translation as delta functions at stimulus onset and convolving these events with a biexponential kernel (rise τ=0.6 s; decay τ=2 s). These parameters were selected to approximate the prolonged calcium transients observed experimentally, which substantially outlasted the intrinsic response kinetics expected for nuclear-localized jGCaMP7f. Separate predictors were constructed for fore-responsive, aft-responsive, and bidirectional (fore plus aft) response profiles. For each ROI, the correlation of trial-averaged ΔF/F waveform with each predictor was computed, and cells with a correlation coefficient exceeding 0.3 were classified as responsive. This threshold was selected empirically because it robustly separated apparent stimulus-responsive cells from baseline fluctuations and was applied uniformly across regions and conditions. Each cell was assigned exclusively to the response category exhibiting its strongest positive correlation. For visualization and population analyses, ΔF/F traces were averaged across all responsive cells within each anatomical region and response category per fish.

### Supervised Learning Model

#### Model construction

We implemented 4 Purkinje cell targets (T) projecting according to matrix *W*_*3*_ to 2 output cells, corresponding to the two orthogonal dimensions in the movement plane (0:180° and 90:270°). We then added 8 Purkinje cells (P) that synapse onto the target cells (T), with each Purkinje cell receiving one IO input tuned to a specific direction in the plane (i.e., 0°, 45°, 90°, 135°, 180°, 225°, 270°, and 315°)^10^, defined in the matrix *A*_*io*_ with dimensionality NxP, where N is the number of Purkinje cells and P is the dimensionality of the output layer. Finally, as we explain later in this section, we added a Purkinje cell input layer representing cerebellar granule cells (g) that sparsely encoded motor commands^118^. Both the olivary input weights, *A*_*io*_, and the target-to-output weights, *W*_*3*_, were held fixed. The model therefore contained two plastic weight matrices, each regulated by the IO: the Purkinje-to-target weights *W*_*2*_, established during development by the ‘wiring rule,’ and the granule-to-Purkinje weights *W*_*1*_, modified during mature learning by the ‘learning rule.’

#### Circuit maturation

During development, granule cells in the model were inactive, and activity in the target cells was drawn from a normal distribution, which in turn generated movement. Because olivary tuning (*A*_*io*_) is fixed and behavioral output is produced by target cells through *W*_*3*_, driving target cell activity randomly and defining IO activity from the resulting output (below) produces correlation between IO cells and target cells with directionally similar sensory tuning and motor function, respectively. This regime reflects an alternate mechanism intended to capture the activity correlations observed in zebrafish when IO neurons and putative Purkinje cell targets are co-activated by sensory experience.

ION activity was defined as taking values 1, –1, or 0 according to a threshold function:

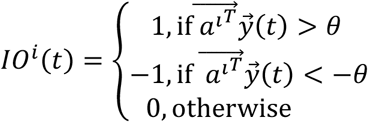

where 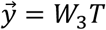, *θ* is a threshold value and *a*^*i*^ is the *i*^*th*^ column of the *A*_io_ matrix. This piecewise function was used to approximate the low frequency activity of IO cells. However, the model performed as robustly when using a simpler linear equation whereby 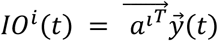. Negative values for IO neurons represent below baseline activity values (Herzfeld *et al*., 2015). Synaptic weight changes between Purkinje and target cells followed a Hebbian rule:

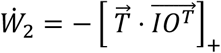

with the additional normalization such that each row of *W*_*2*_ was divided by its maximum. Effectively, this learning rule drove positive correlation of the tuning of the IO and target cells linked by Purkinje cells in order to set 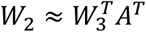 (see **Appendix Part B**). This Hebbian developmental rule is compatible with our experimental finding that maturation of Purkinje cell projections requires activation of the IO and the acquired target, and with the established role of correlated activity in wiring developing circuits^59,119^.

#### Mature activity and learning

After an initial phase of circuit development, activity in the target cells became driven by Purkinje cell input which in turn was driven by granule cell input such that,

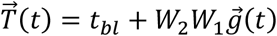

where *tbl* is the baseline activity of target cells to ensure their firing remained positive and could vary above and below baseline values. 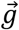 is the granule cell activity where the 2xN_g_ matrix, *A*_*g*_, defines the direction of intended movement in which each granule cell is active. N_g_ is the number of granule cells. Then, given a movement instruction 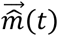, and defining 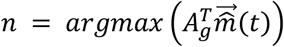 :

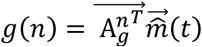

and all other granule cells = 0. (For example, if N_g_=180, each granule cell was only active for instructions within 1 degree of its peak tuning response).

We adopted the canonical error-based formulation in which mature IO activity encodes the difference between desired and actual output^47,120^. Error in the movement was defined as:

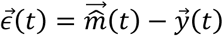

and IO activity post development was:

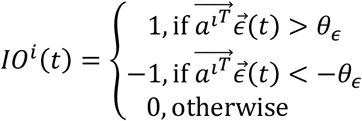

Learning from error at the granule-Purkinje cell synapses followed an anti-Hebbian rule:

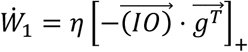

Effectively, this learning rule produces anti-correlated tuning of the granule and IO cell inputs to a Purkinje cell.

#### Effective weight

To measure the effect of the motor command to propagate through the network and produce both a matching movement in the desired direction and no movement in the direction orthogonal to the desired direction (the null direction), we calculated the following:

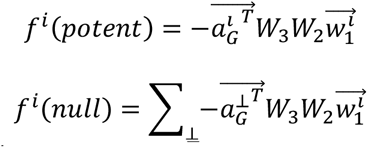

where 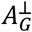 are the directions orthogonal to 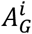.

#### Measuring alignment

As we show in the **Appendix**, the key parameter for this network to learn any task is that both eigen values of the symmetrized matrix, *B*, are positive. In our simulations, we used the Frobenius inner product of *W*_2_ with 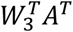 as a measure for alignment as this value provides the normalized sum of the eigen values and is therefore a good proxy of alignment.

### Statistical Analysis

All statistical analyses were performed in MATLAB. For two-group comparisons of continuous measures (*e*.*g*., axon length, bouton width, bouton density), two-tailed Student’s t-tests were used. For comparisons across more than two groups with a single factor (*e*.*g*., stimulation age, temperature), one-way ANOVA was used followed by Bonferroni-corrected post-hoc tests for pairwise comparisons. For experiments with two crossed factors (*e*.*g*., CM × lesion condition), two-way ANOVA was used to assess main effects and interactions. For non-parametric paired comparisons (namely, within-brain laterality of ION lesion completeness), the Wilcoxon signed-rank test was used. Behavioral data in which repeated measurements were obtained from groups of fish across rounds were analyzed with linear mixed-effects models to account for the nested structure of repeated observations within groups, with fixed effects of treatment condition and stimulus type, and significance was assessed by F-tests on fixed effects. For binary outcomes (*e*.*g*., presence of MON innervation), logistic regression was used (generalized linear model with binomial distribution and logit link function, implemented with fitglm). Predictors were included as main effects and their interaction; significance of each term was assessed by likelihood-ratio χ^2^ tests comparing the full model to a reduced model lacking that term. Sample sizes (*n*) refer to individual brains for anatomical measures and to groups of simultaneously evaluated fish for behavioral assays. Where data are plotted as bar graphs, values represent mean ± SEM unless otherwise noted. Significance thresholds are denoted as ^*^ *p*<0.05, ^**^ *p*<0.01, ^***^ *p*<0.001, ^****^ *p*<0.0001; *ns*, not significant, *p*>0.05.

## Data availability

Raw calcium imaging data and simulation results as well as summary anatomical and behavioral data will be posted to Dryad upon publication and are also available from the authors upon request.

## Code availability

Analysis and simulation code will be posted to Dryad upon publication and are also available from the authors upon request.

## Author contributions

Imaging: RJM, KM, TF, JL, KW, DEE; Transgenics: RW, SN, MA, MH; Animal behavior: RM, DEE; Mathematical analysis: BW & SM; Computational modeling: BW, SM, DEE; Data analysis: RJM, KM, SJ, JM, CK, JD, NM, RW, EAS, DEE; Visualization: RJM, SM, BW, TF, DEE; Conceptualization: RJM, SM, BW, DEE; Funding acquisition: JH, KWE, DEE; Supervision: RJM, SM, DEE; Writing: RJM, SM, BW, DEE.

## Competing interests

TF is the owner of Forward Scientific LLC, which contributed to the design and construction of the custom voice coil drive amplifier used for the μSled device in this study, and for generating CAD drawings of the device.

## Acknowledgments

The authors would like to thank David Herzfeld, Eric Lang, Larry Abbott, Nate Sawtell, Sascha du Lac, Nimish Pujara, and Betsy Quinlan for helpful advice on experimentation and interpretation. Research was supported by the National Institute of General Medical Sciences of the National Institutes of Health under award number R35GM146885 to DEE, Beckman Foundation support for the Beckman Center for Advanced Light Microscopy at the Morgridge Institute for Research, the Simons Foundation, Kavli Foundation, Swartz Foundation, and the Gatsby Charitable Foundation.

Supplementary information is available for this paper.

## Supplemental Figures

**Supplemental Figure S1.**
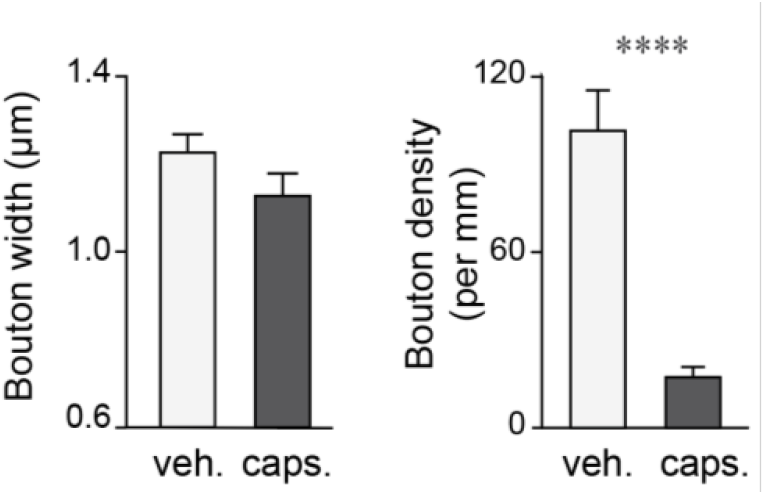
Immature bouton properties of Purkinje cell axons following direct stimulation. Median bouton width (*left*) and bouton density (*right*) along Purkinje cell axons in the flow nucleus (mean ± SEM) in larvae with and without 1 µM capsaicin exposure (width: Student’s t-test, t(13) = 1.56, not significant; density: t-test, t(10) = 2.49, ^*^*p*< .05).

**Supplemental Figure S2.**
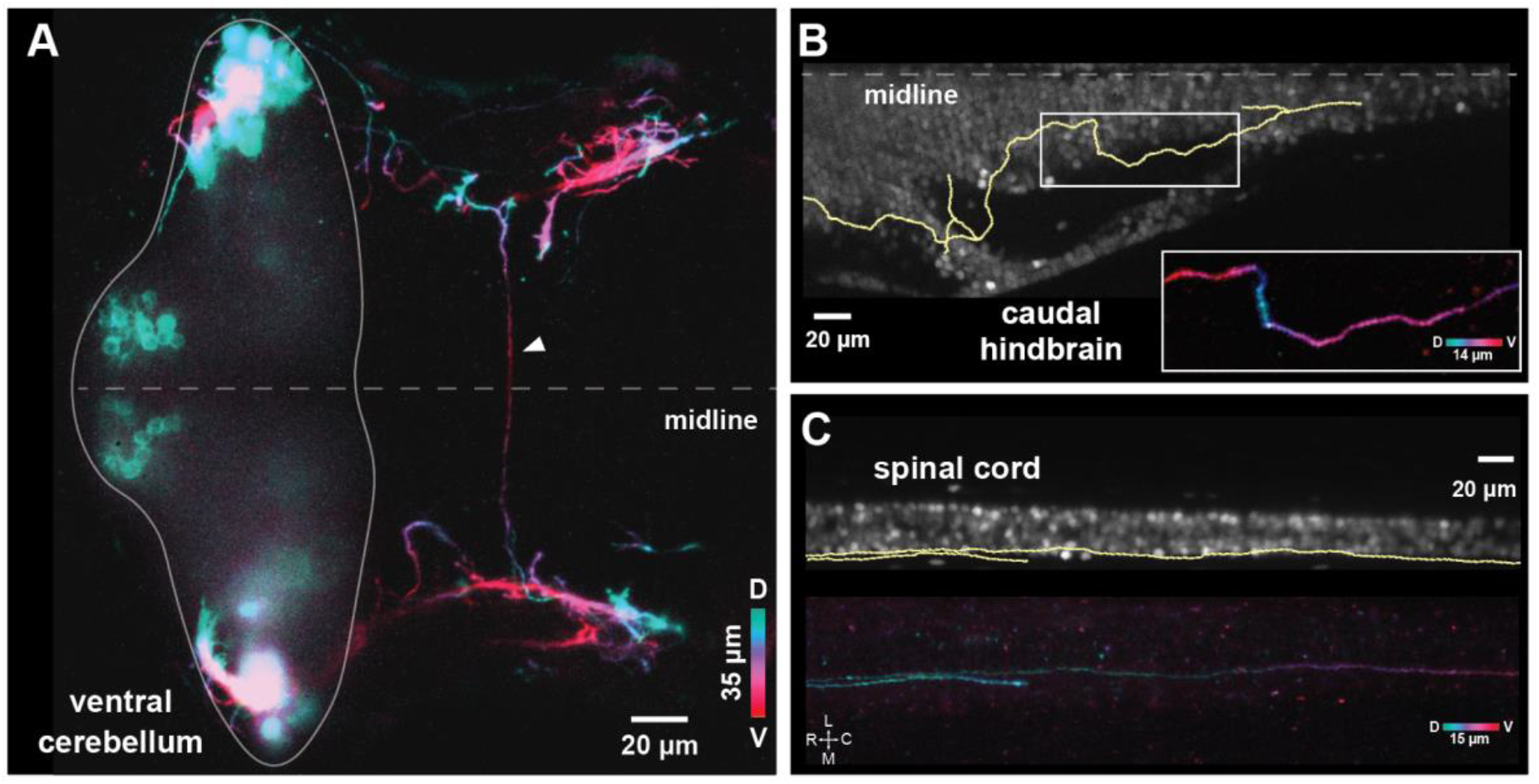
Purkinje cell stimulation induces ectopic projections. (**A,B,C**) Representative micrographs of Purkinje cell axons exhibiting atypical projections following 48-hr exposure to 1 µM capsaicin (12 days, *Tg(aldoca:TRPV1-tagRFP; elav3l:H2B-jGCaMP7f*)), including (*A*) commissural projections, (*B*), projections into the caudal hindbrain, and (*C*) projections into the spinal cord.

**Supplemental Figure S3.**
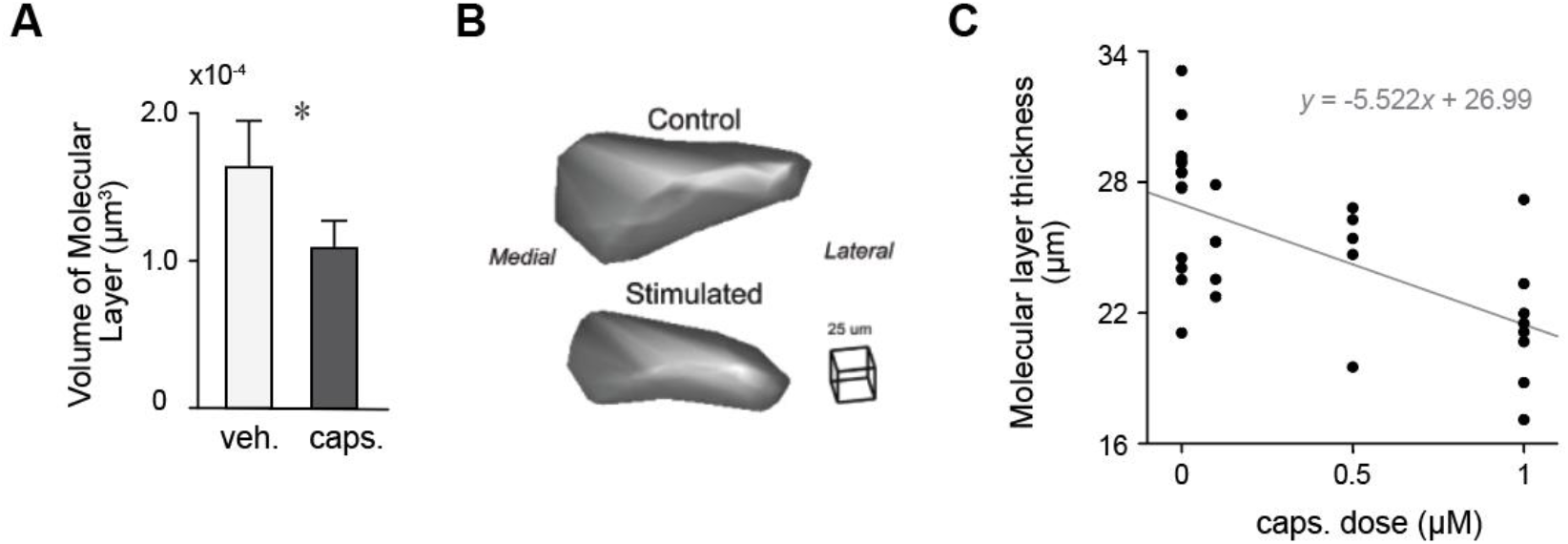
Purkinje cell stimulation reduces cerebellar molecular layer size. (**A**) Average molecular layer volume is reduced by 33.1% in larvae exposed to 1 μM capsaicin for 48 hr compared to controls (8 days, *Tg(aldoca:TRPV1-tagRFP; elavl3:H2B-jGCaMP7f)*, Student’s t-test, t(8) = 3.39, ^*^*p*< 0.05). (**B**) Representative 3D renderings of molecular layers from two samples (right cerebellar hemisphere). (**C**) Molecular layer thickness decreases with increasing capsaicin dose (0, 0.1, 0.5, 1 µM; 48 hr). Thickness was measured at the maximal width of the layer; gray line indicates linear regression fit.

**Supplemental Figure S4.**
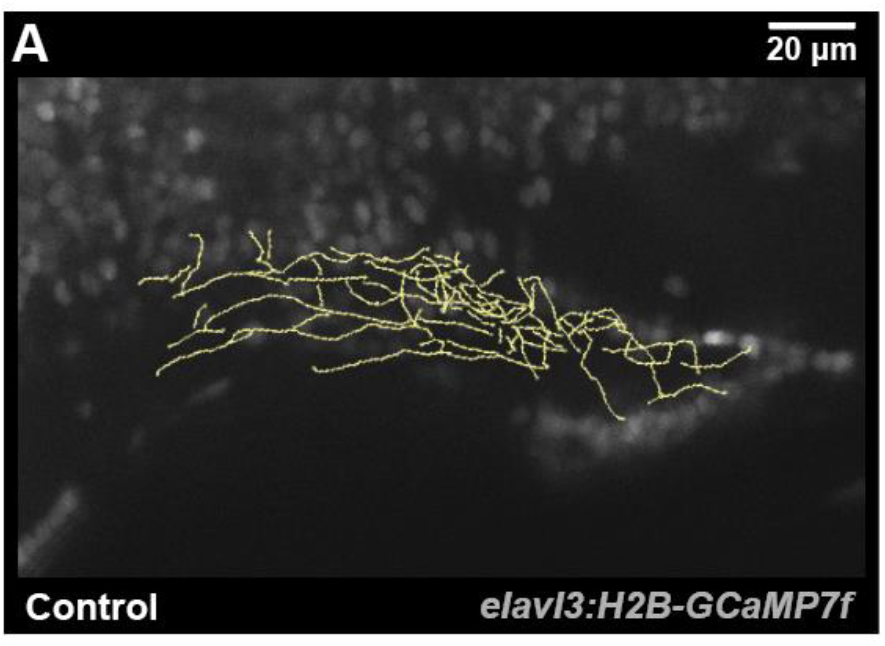
Representative Purkinje cell projections to vestibular nuclei. Horizontal projection of traced Purkinje cell axons within the vestibular nuclei (yellow) in one hemisphere of a representative fish, overlaid on a single corresponding micrograph of neuronal nuclei (12 days, *Tg(aldoca:TRPV1-tagRFP (not shown); elavl3:H2B-jGCaMP7f)*).

**Supplemental Figure S5.**
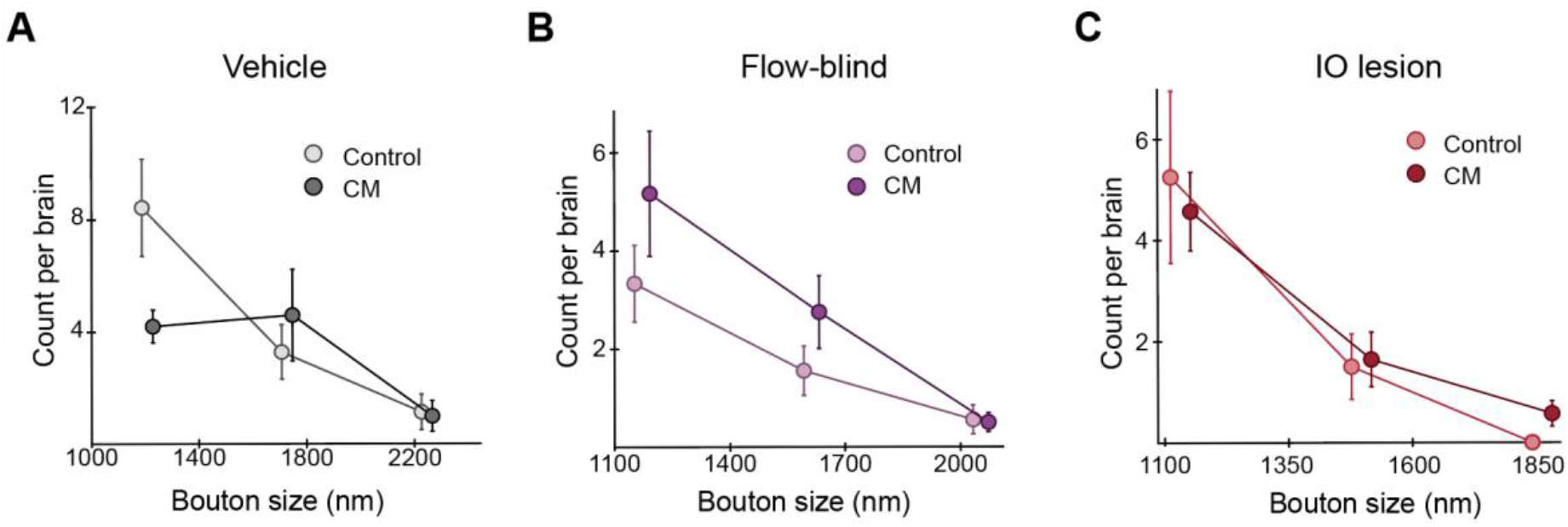
Distribution of bouton sizes after CM. Histograms of bouton size for Purkinje cell axons in the flow nucleus at 12 days across three bins for larvae raised with CM and sibling controls (mean ± SEM across individuals, n=8).

**Supplemental Figure S6.**
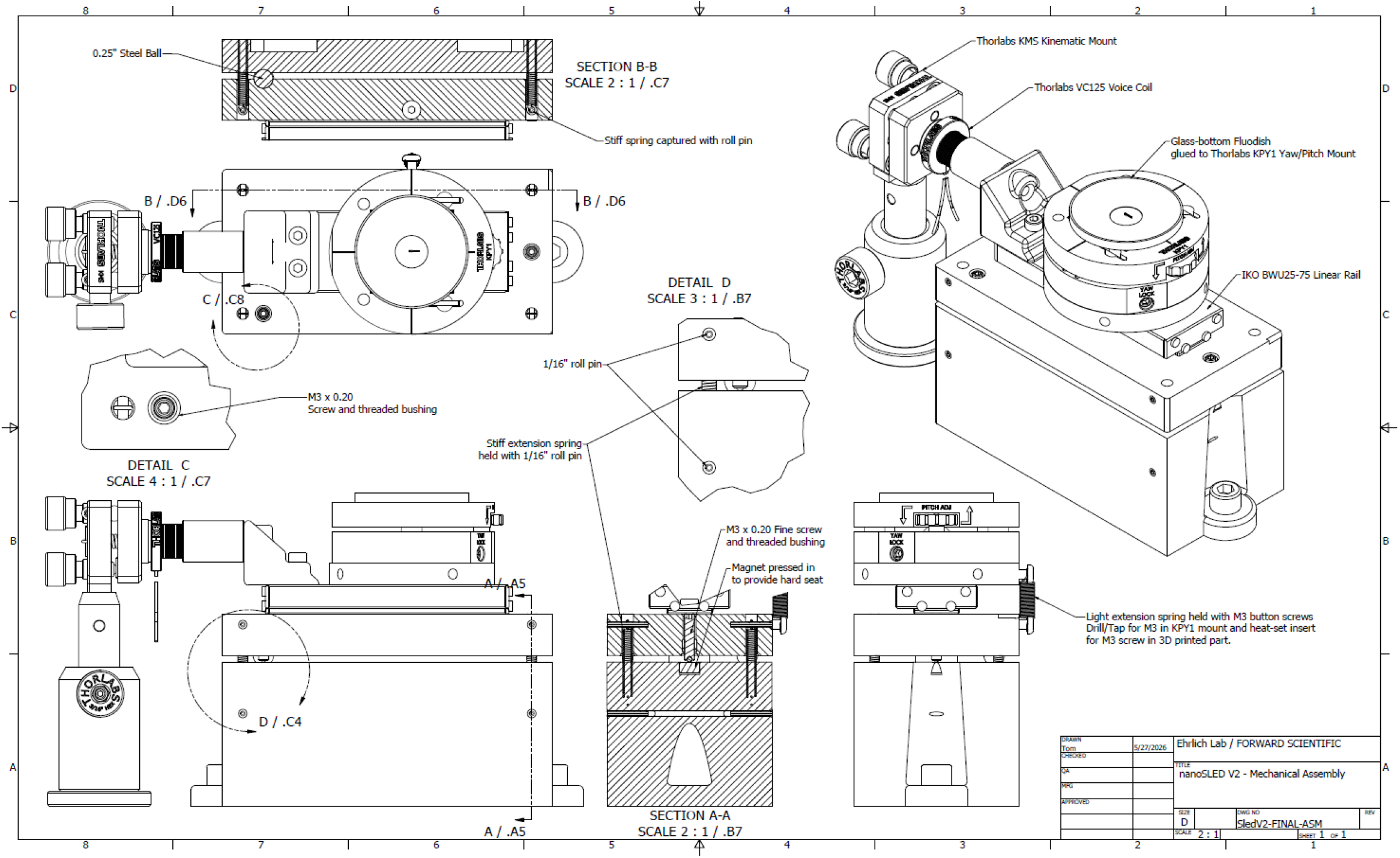
Computer-assisted design drawing of μSled constructions. See *Bill of Materials* in Table 1 for parts.

**Supplemental Figure S7.**
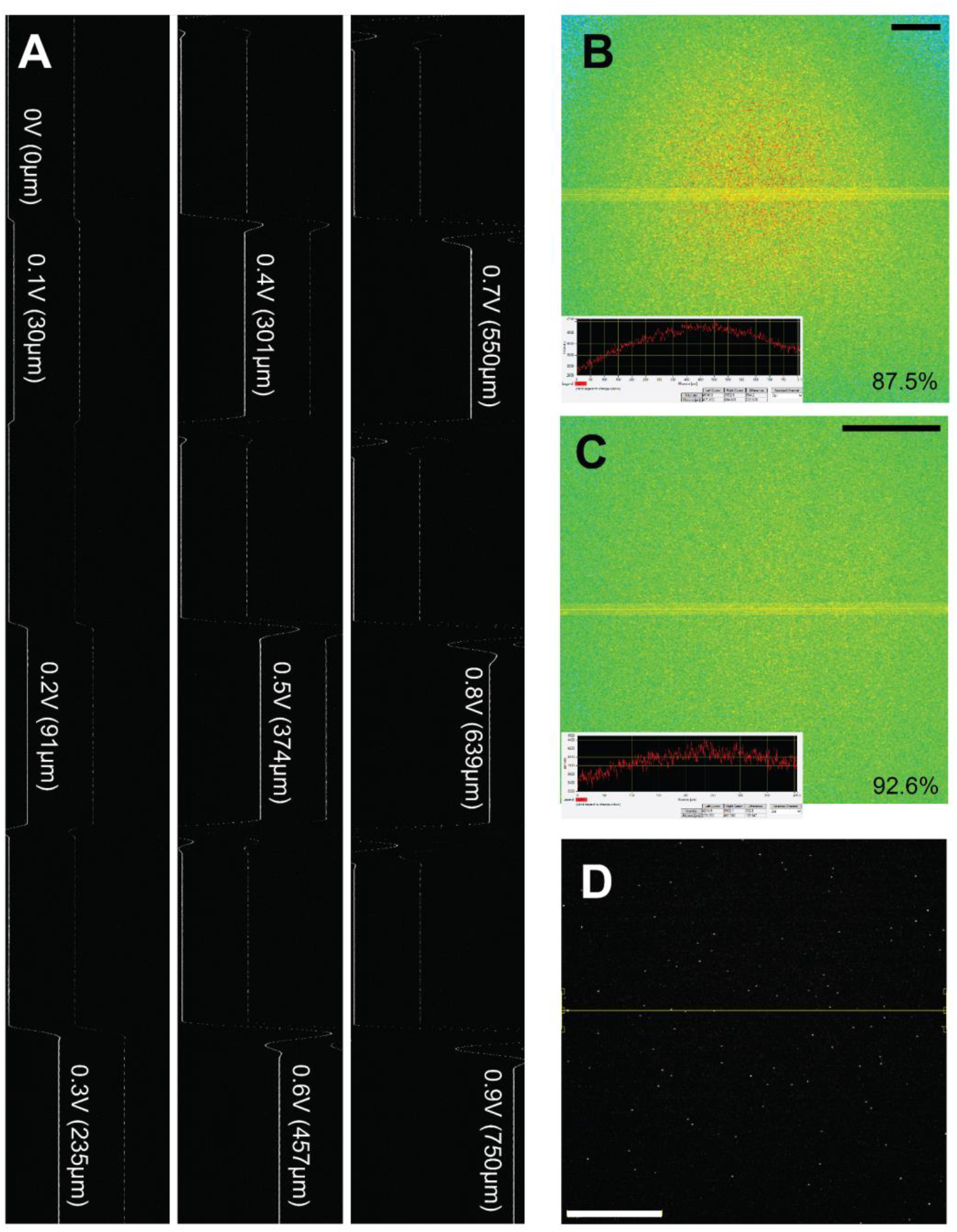
μSled motion characterization. **(A)** Two-photon line-scan imaging of fluorescent beads on the uSled during a step function. Steps were incremented by 0.1V drive and held for 1 sec before a reset to the 0V center position. **(B,C)** Two-photon image of a fluorescent slide showing field uniformity across the entire 812um field (*B*, 1x zoom, scale bar 100mm) and at 2x zoom (*C*, scale bar 100mm). **(D)** Whole-field image of 1μm beads showing position of line scan in panel *A* (scale bar 100mm).

**Supplemental Figure S8.**
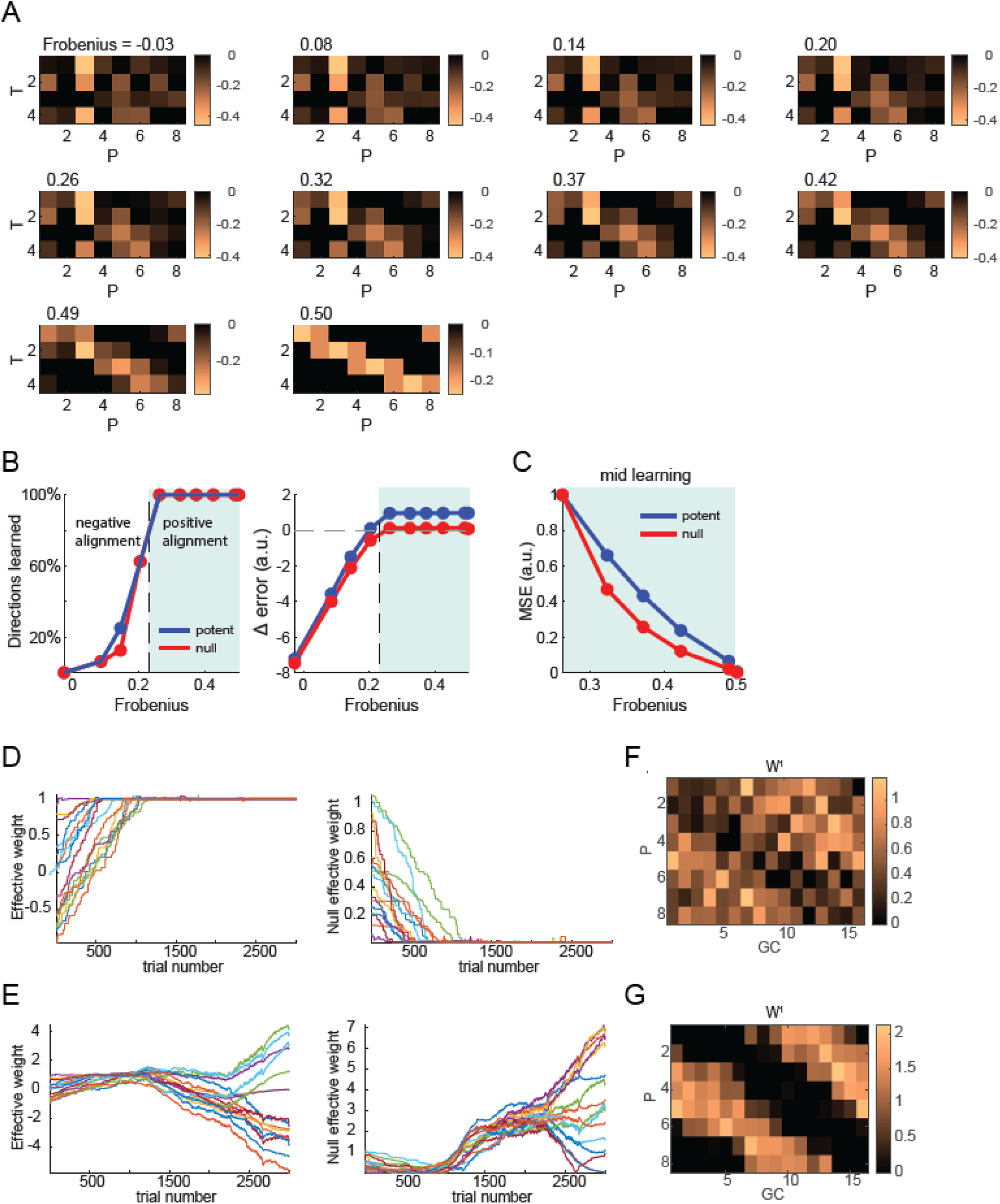
Positive alignment of instruction and downstream function is required for learning. (**A**) All connectivity patterns between Purkinje (P) and target cells (T) used in the simulations. The title denotes the Frobenius value. (**B**) Effect of alignment on performance of learning in the instructed (potent) and orthogonal (null) direction. The total error reported in the paper is the sum of these errors. For networks with positive alignment (shaded area), learning converges such that all instructed directions are learned (*left*) and error relative to initial error decreases (*right*). The vertical dashed line denotes the Frobenius value at which both eigenvalues become positive. (**C**) Learning is faster with greater alignment, resulting in faster reduction of the mean squared error (MSE) in both the potent and null directions. (**D, E**) For the aligned *W*_*2*_ (*D*) learning in all directions (where each line represents one of 16 directions) converges such that in the potent direction the effective weight (defined in the *Methods*) converges to one (*left*) and converges to zero in the null direction (*right*). For the misaligned *W*_*2*_ (*E*) learning diverges in all directions. (**F, G**) *W*_*1*_ at the end of the simulation for the aligned (*F*) and the misaligned (*G*) *W*_*2*._

**Supplemental Figure S9.**
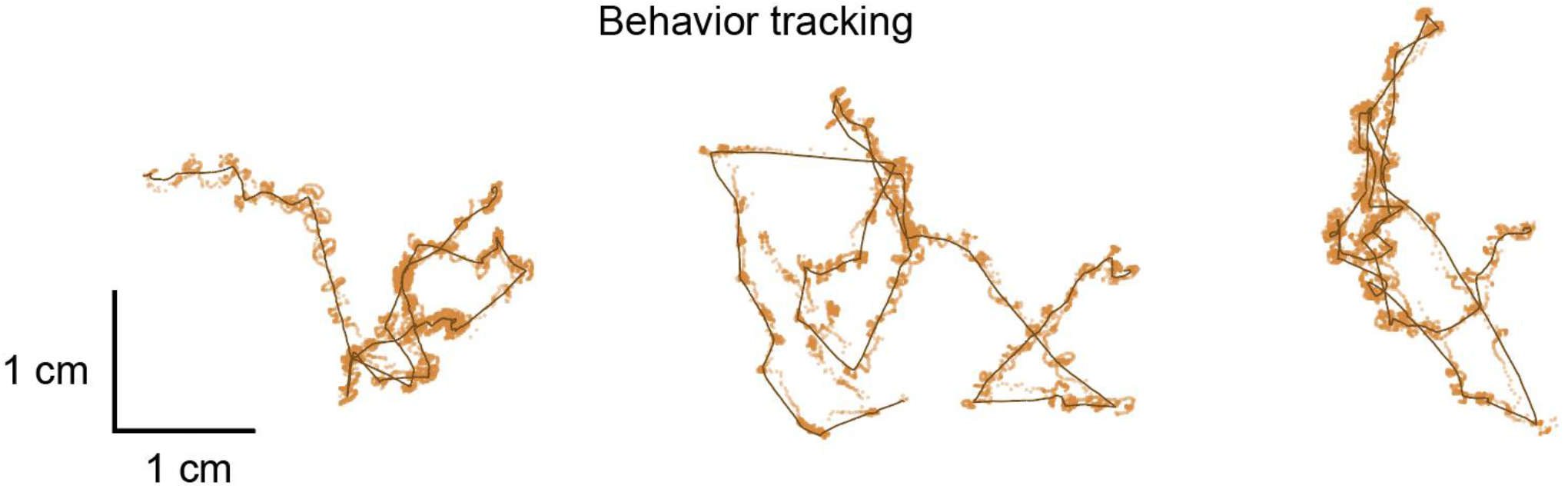
Example swimming trajectories from flow assay. Representative traces from individually tracked 12 days fish showing raw position data (orange dots) and low-pass filtered trajectories (lines).

**Table 1.**
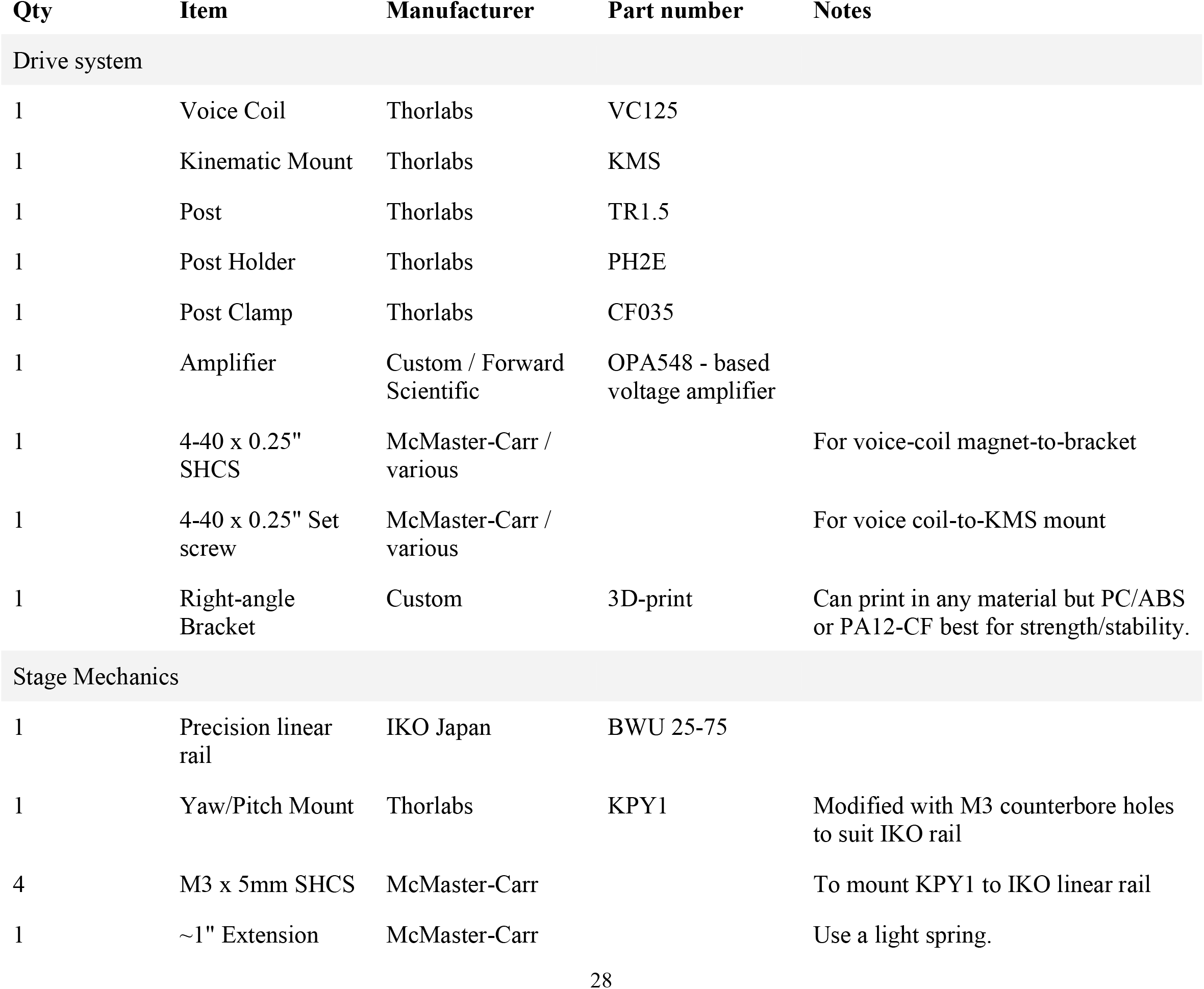
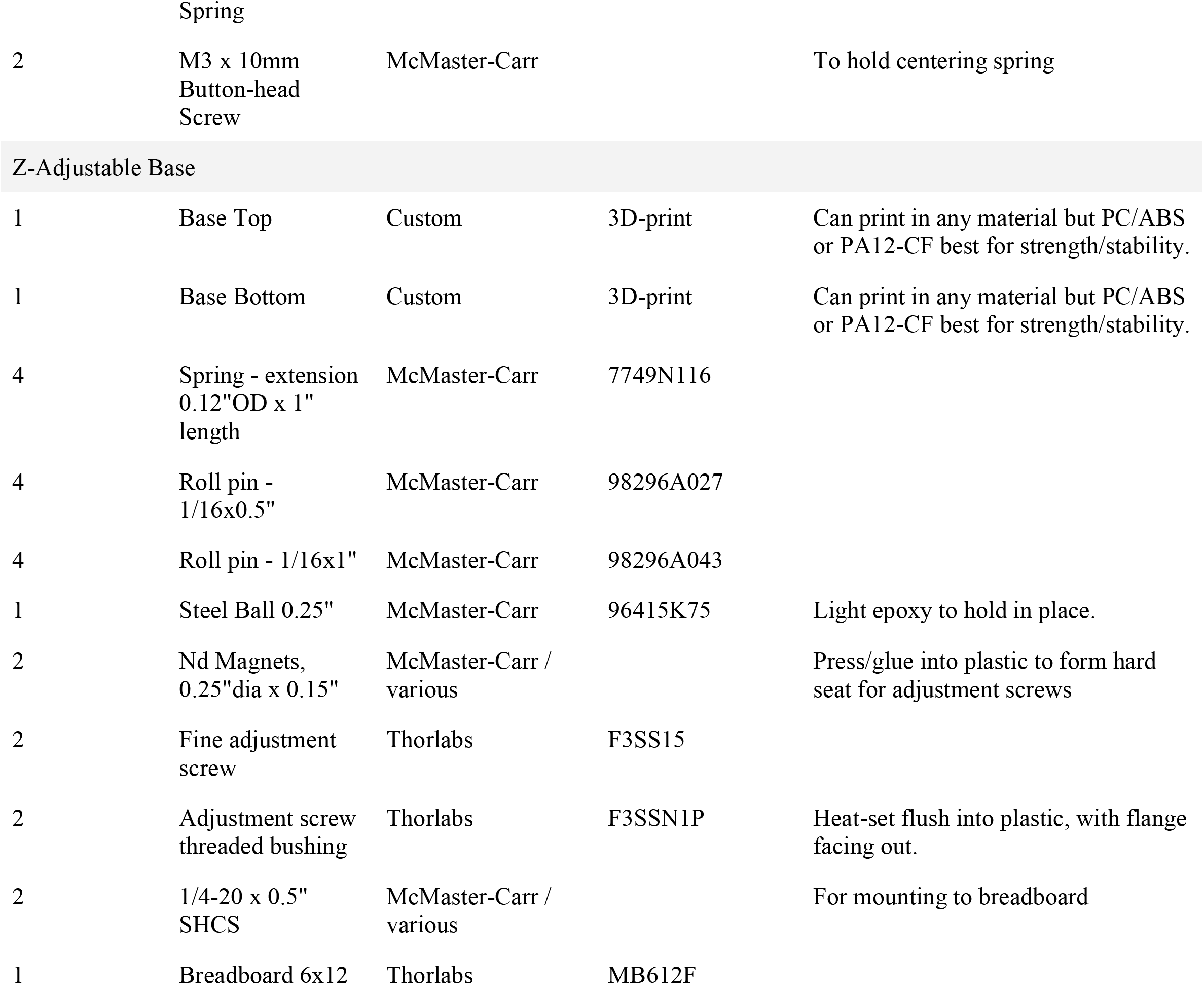
Bill of Materials for μSled.

## Appendix

We consider the four-layer network (Granule cells → Purkinje cells → Target nuclei → Output)

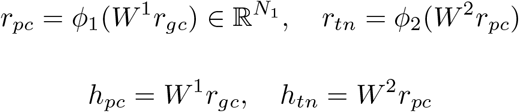

where 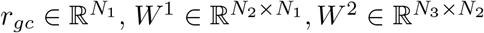 and 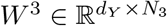, and we assume that *W*^3^ is fixed. For convenience, we denote the neuron activity in each layer as

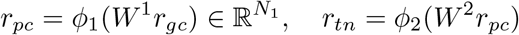

The training data is 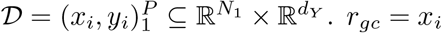 and we sample one data point (*x*_*i*_, *y*_*i*_) during each training step (single-batch). The learning of *W* ^1^ at each iteration is driven by climbing fiber input to the Purkinje cells (*r*_*cf*_) and follows a modified delta rule in which the error is weighted through the matrix 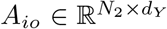,

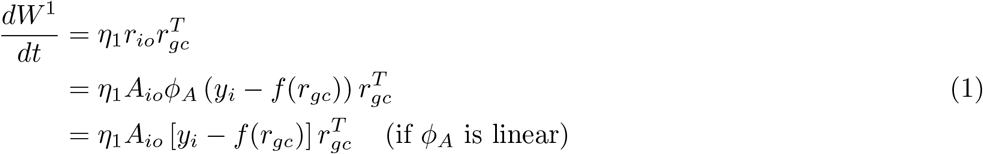

The loss function is then,

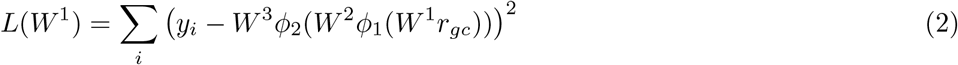

The loss change following learning is,

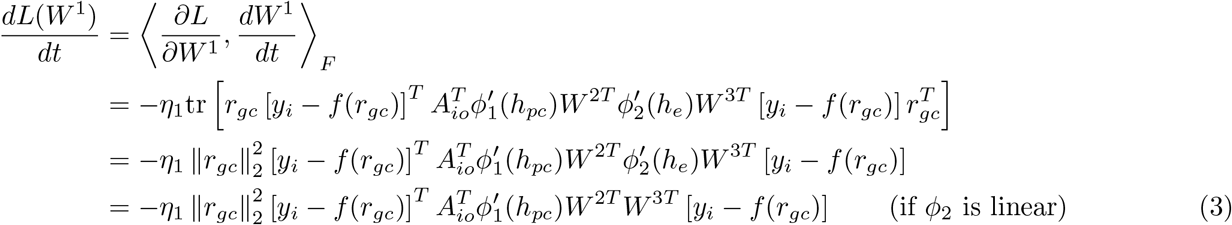

Note that 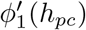 and 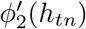 represent diagonal matrices whose elements are given by the vector 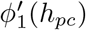 and the vector 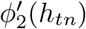 respectively.

The above loss change has the form ™*x*^*T*^ *Mx*, and it is negative for all *x* if the matrix

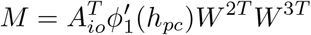

is positive definite, guaranteeing a decrease of the loss function.

1. Thus, based on this form, we derive the following *sufficient* (results 1&2) and *necessary* (results 3&4) conditions:
2. For a linear function or ReLU *ϕ*_1_ that operates in the linear regime, 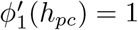 (and since Purkinje cells have high baseline firing rate, they are assumed to operate in approximately a linear regime). Then,

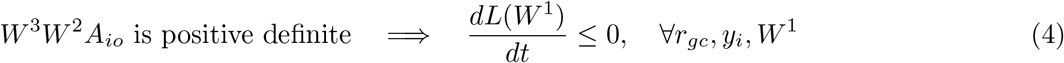
3. For monotonically non-decreasing nonlinearity *ϕ*_1_ (saturating or non-saturating), 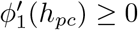. Then,

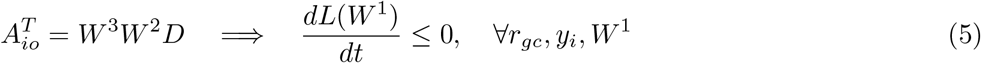

where *D* is a diagonal matrix of all positive values. Defining *W* ≡ *W*^3^*W*^2^, we have 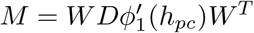, since 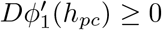, *M* is positive semidefinite.
4. For a linear or any monotonically increasing nonlinearity *ϕ*_1_, 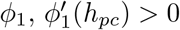. Then,

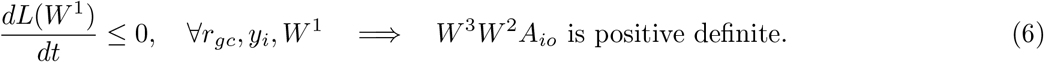

This means that for some tasks *W*^3^*W*^2^*A*_*io*_ must be positive definite for the loss to decrease (see Proof below), and that the opposite (i.e. making *W*^3^*W*^2^*A*_*io*_ not positive semidefinite) is not useful for any task (shown in 1&2). Note that this is a sufficient (result 1) and necessary condition for loss to decrease with linear *ϕ*. **Proof:** We can choose *W*_1_ such that *h*_*pc*_’s are all equal (for example by making all rows of *W*_1_ equal), making *ϕ*^*′*^(*h*_*pc*_) behaves like a positive scalar and thus decrease in loss is equivalent to

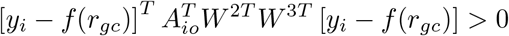

Since *y*_*i*_ and *r*_*gc*_ can be arbitrary, it means that *W*^3^*W*^2^*A*_*io*_ is positive definite.
5. For saturating monotonically increasing nonlinearity *ϕ*_1_, 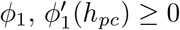 Then,

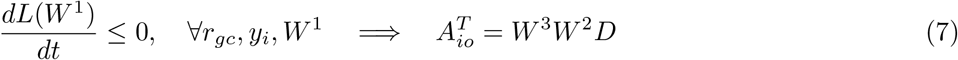

where *D* is a diagonal matrix of all positive values. This means that for some tasks 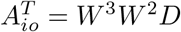 for the loss to decrease (see Proof below), and that the opposite (i.e. *A*_*io*_ deviating from *W*^3^*W*^2^*D*) is not useful for any task (shown in 2). Note that this is a sufficient (result 2) and necessary condition for loss to decrease with saturating nonlinearity *ϕ*. **Proof:** We can choose *W*_1_ and *x*_*i*_ such that *ϕ*^*′*^(*h*_*pc*_) equal to the *i*th standard basis (by making all but one Purkinje cell in the saturated regime) and the matrix *M* becomes

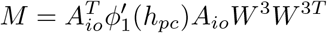

This rank-one matrix is positive semidefinite iff

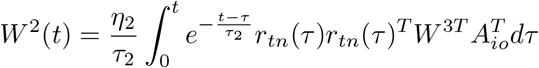

(where *d*_*i*_ is a positive scalar). This equality is equivalent to

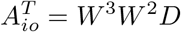

B. Here we show how a *W*^2^ matrix obtained via Hebbian plasticity during development may satisfy the alignment condition. Suppose that during development, the efference layer has spontaneous activity *r*_*tn*_(*t*), which drives the activity in the PC layer through ION connectivity: *r*_*io*_ = *A*_*io*_*W*^3^*r*_*tn*_. Then *W*^2^ undergoes Hebbian plasticity,

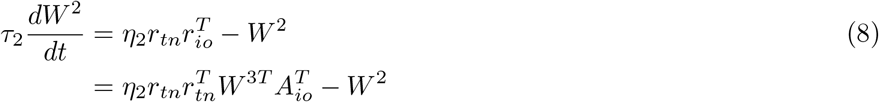

Solving for *W*^2^, we get

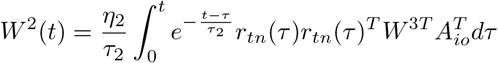

If *τ*_2_ ≫ 1, the steady state weight is well approximated by its mean,

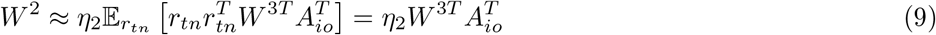

Thus, we have

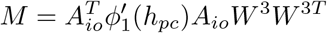

If *W*^3^ is random and *N*_3_ ≫ 1 or if *W*^3^ has orthogonal rows of equal norm (i.e. the constructed push/pull effector system we use in the model) then *W*^3^*W* ^3*T*^ → *σ*_3_*I*_*d*_ and thus,

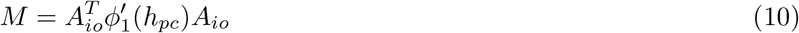

making *M* positive definite (given 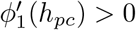)!

Similarly, if *A*_*io*_ has orthogonal columns of equal norm (as in our simulated model) and *ϕ*_1_ is linear, *M* = *W*^3^*W*^3*T*^ - a positive definite matrix!

